# Overexpression of *PtaHDG11* enhances drought tolerance and suppresses trichome formation in *Populus tremula* × *Populus alba*

**DOI:** 10.64898/2026.01.12.699028

**Authors:** Alexander Fendel, Matthias Fladung, Tobias Bruegmann

## Abstract

Extended drought periods are increasingly impacting the growth and survival of forest trees in Central Europe, underscoring the need for innovative approaches to adapt trees to drought. Understanding the functions of genes involved in drought tolerance can contribute to the development of effective strategies to improve drought resilience. Although the homeobox-leucine zipper protein HDG11 has been identified as a promising target for enhancing drought stress tolerance in plants, the functional role of the native *HDG11* gene in tree species remained uncharacterized. To address this gap, we identified the homolog of the *AtHDG11* gene in the poplar hybrid *Populus tremula* × *Populus alba*, clone INRA 717-1B4, and subsequently transformed the *PtaHDG11* gene into this clone for constitutive overexpression. Drought stress experiments revealed significantly enhanced drought stress tolerance in *PtaHDG11*-overexpressing poplars compared with the wild type. This improved tolerance was characterized by higher leaf relative water content, reduced leaf shedding, lower malondialdehyde accumulation and higher expression of antioxidant-related genes, including *SOD* and *CAT*. Improved tolerance resulted in significantly increased dry biomass after the recovery phase. Notably, the transgenic poplars exhibited altered basal expression of the cell wall-related genes *PtaXTH32*-*like* and *PtaEXPA15*-*like* under control conditions, indicating a modified cell wall-associated regulatory state before stress exposure. In addition, *PtaHDG11* overexpression was associated with a glabrous phenotype lacking trichomes, pointing to functions of *PtaHDG11* beyond drought stress responses in a woody perennial. Together, these findings suggest that *PtaHDG11* can contribute to improving drought resilience in trees, with implications for sustainable forest management and breeding programs.

## 1 Introduction

The genus *Populus* comprises economically important tree species that are distributed in temperate regions worldwide. Owing to their rapid growth, modest environmental requirements, wood quality, and ease of propagation, poplars are cultivated in short-rotation coppices. Poplars are often utilized in various industrial applications, such as the production of raw materials for the timber industry and paper manufacturing (Komán et al., 2023), as well as the production of biofuels and biorefineries, contributing to renewable energy sources (An et al., 2021). Furthermore, the use of poplar for genetic functional studies yields important information for refined forest tree breeding and the potential direct use of genetically modified poplar, where permitted. In this context, investigating the mechanisms underlying drought tolerance in poplars is essential, as this knowledge can be applied to improve the resilience of forest ecosystems, which are under increasing threat due to climate change. In particular, over the past eight years, European forests have been increasingly vulnerable to weather extremes, with notable increases in the frequency and intensity of summer droughts (Schuldt et al., 2020; Bastos et al., 2021). Such drought periods are predicted to become even more frequent and severe in the coming years, with climate models forecasting a significant increase in drought intensity and duration across European forests (Knutzen et al., 2025). Therefore, the breeding of drought-tolerant trees, including poplars, has become a challenging demand (Zhao et al., 2023).

Drought stress tolerance refers to a plant’s ability to maintain the potential for optimal physiological functions, growth, and survival under water-limited conditions (Fang and Xiong, 2015). This encompasses both the plant’s capacity to mitigate the negative impacts of drought through various morphological, physiological, molecular physiological, and cellular adjustments during stress and its subsequent ability to recover efficiently and resume productive growth (fitness) once water availability is restored (Polle et al., 2018). Crucially, the long-term productivity and reproductive success of a plant following a drought event are key indicators of true tolerance, extending beyond mere survival during the stress period.

Drought stress leads to the formation of cellular reactive oxygen species (ROS), which are generated through the disruption of cellular homeostasis, such as the inhibition of photosynthetic electron transport. The accumulation of ROS can also trigger a range of signaling pathways, including those involved in the stress response and programmed cell death (Apel and Hirt, 2004). The detrimental effects of ROS can be mitigated by the action of antioxidative enzymes, such as superoxide dismutase (SOD) or catalase (CAT). SOD and CAT are essential components of the plant antioxidant defense system and play crucial roles in detoxifying ROS. SOD acts as the primary defense mechanism against ROS, catalyzing the conversion of superoxide anions to hydrogen peroxide and molecular oxygen. Additionally, CAT further detoxifies hydrogen peroxide by converting it into water and oxygen, thereby protecting plant cells from oxidative damage (Leung, 2018). Molecular physiological responses to drought include the accumulation of proline, an amino acid with osmoprotectant properties, which has been observed in various plant species, including several hybrid poplar genotypes and *Populus euphratica* (Chen et al., 2003; Barchet et al., 2014).

The key players in those stress adaptive mechanisms are transcription factors (TFs), which can regulate the expression of genes involved in stress responses (Joshi et al., 2016; Manna et al., 2021). The homeodomain-leucine zipper (HD-ZIP) protein family plays a central role in plant development, including processes such as organogenesis and leaf development (Roodbarkelari and Groot, 2017; Tu et al., 2022). For example, HD-ZIP IV members bind HD-ZIP-cis elements that are frequently present in the promoters of expansin, extensin, or xyloglucan-endo-transglycosylase/hydrolase (*XTH*) genes, thereby regulating cell-wall extensibility (Xu et al., 2014). HD-ZIP TFs are also involved in stress responses. In particular, the *HOMEODOMAIN GLABROUS 11* (*HDG11*) gene, which belongs to HD-ZIP subfamily IV, has emerged as a promising target for enhancing drought tolerance. At present, functional studies of *HDG11* have focused almost exclusively on *AtHDG11* from the model plant species *Arabidopsis thaliana.* The increased expression of *AtHDG11* in *A. thaliana* and *Nicotiana tabacum* significantly enhances drought tolerance by reducing leaf stomatal density, increasing root system development, increasing proline and abscisic acid accumulation, and, in general, activating genes involved in stress response pathways (Yu et al., 2008). Similarly, the overexpression of *AtHDG11* in tall fescue (*Festuca arundinacea*) improved drought and salt stress tolerance via increased ROS-scavenging ability (Cao et al., 2009). Similar results, achieved through heterologous overexpression of *AtHDG11*, have been reported for sweet potato (*Ipomoea batatas*; Ruan et al., 2012), rice (*Oryza sativa*; Yu et al., 2013), Chinese kale (*Brassica oleracea*; Zhu et al., 2016), and upland cotton (*Gossypium hirsutum*; Yu et al., 2016). In addition, Yu et al. (2016) reported that *AtHDG11* overexpression in Chinese white poplar (*Populus* × *tomentosa*) improved drought and salt stress tolerance, resulting in a more developed root system, reduced leaf stomatal density, and increased stomatal and leaf epidermis cell size.

While the functions of the *AtHDG11* gene have been widely studied in herbaceous plants and has also been investigated in *Populus* × *tomentosa*, detailed analyses of the associated physiological and molecular responses in tree species are still lacking. Furthermore, consequences following enhancing the activity of the endogenous *HDG11* homologs in trees remained unexplored. Here, we address this knowledge gap by investigating the role of *PtaHDG11* in drought tolerance in the poplar hybrid *P. tremula* × *P. alba* using morphological, physiological, and molecular analyses. Notably, despite drought stress tolerance being a complex trait involving multiple genes, the overexpression of a single gene copy from the *P. alba* allele (gene number *PtXaAlbH.15G026700*) in the poplar hybrid clone significantly improved drought stress tolerance under controlled conditions, making it a promising target for further research and contributing to the development of forest stands with increased drought resilience. Additionally, we also revealed that this gene is involved in trichome formation.

## 2 Materials and Methods

### 2.1 Identification and bioinformatic analysis of candidate gene

To identify the *PtaHDG11* gene in *P. tremula* × *P. alba*, we utilized the protein sequence of *At*HDG11 from *A. thaliana* (*AT1G73360*) as the query sequence. A BLAST search was performed against the genomes of *P. tremula* × *P. alba* HAP1 v5.1 and HAP2 v5.1, which were accessed through the Phytozome database (Goodstein et al., 2012). The protein sequences of *A. thaliana*, both *P. tremula* × *P. alba* alleles, and *Populus trichocarpa* were retrieved from the Phytozome database and aligned using the Multiple Sequence Alignment by CLUSTALW tool (https://www.genome.jp/tools-bin/clustalw) and subsequently visually adjusted via Jalview (Waterhouse et al., 2009). Protein domains were identified via the SMART web resource (Letunic et al., 2021). A phylogenetic tree was constructed via the Maximum Likelihood Method (bootstrap with 1000 replications) using the MEGA11 program (Tamura et al., 2021). The tree included all genes of the *A. thaliana* HD-ZIP IV class (as described in Nakamura et al. (2006)) and the poplars’ *AtHDG11* homologous genes.

### 2.2 Plant materials and *in vitro* growth conditions

The study was conducted with the poplar hybrid clone INRA 717-1B4 (*P. tremula* × *P. alba*; Leple et al. (1992)). The maintenance and propagation of the wild type (WT) and the subsequently produced transformation lines were performed *in vitro* on solid McCown Woody Plant Medium supplemented with vitamins (WPM; supplied by Duchefa, Haarlem, The Netherlands), 2% (w/v) sucrose, and 0.6% (w/v) agar at a pH of 5.8. The *in vitro* cultures were maintained in Magenta vessels under controlled conditions at 23 °C with approximately 50% relative humidity (RH). Continuous 24-hour light was provided by LED lamps, which delivered a photosynthetic photon flux density of approximately 170 µmol m^−^² s^−1^.

### 2.3 Transformation vector and candidate gene transformation

The binary vector for the constitutive overexpression of *PtaHDG11* was obtained from DNA Cloning Service (Hamburg, Germany; Fig. S1). The T-DNA region of the vector contained the 35S::*nptIIo* selection marker gene, as well as the coding sequence (CDS) of *PtaHDG11*, which was driven by an *AtUBQ10* promoter. The recombinant plasmid was introduced into the *Agrobacterium tumefaciens* strain GV3101-pMP90RK and subsequently transformed into the poplar WT via a modified leaf disc transformation method (Bruegmann et al., 2019). Successfully transformed poplars were regenerated on WPM supplemented with cefotaxime (500 mg/L) to eliminate residual *A. tumefaciens*, kanamycin (50 mg/L) for the selection of successful transformants, and Pluronic F-68 (100 mg/L) and thidiazuron (0.0022 mg/L) to promote efficient regeneration of the plantlets.

### 2.4 Verification of genetic transformation in putative poplar lines

After transformation, nine regenerants were obtained, of which only three lines survived the antibiotic selection process. Genomic DNA was isolated from the regenerated transgenic poplar lines HDG11-1, HDG11-2, and HDG11-3 via the method described by Bruegmann et al. (2022). The putative transgenic lines were identified via standard PCRs targeting the *nptIIo* and *PtaHDG11* genes with primers located on the promoter and terminator of the cassettes. Southern blot analysis was subsequently performed to determine the number of integrated T-DNA sequences in the putative overexpression lines (Fladung and Ahuja, 1995). Additionally, PCRs targeting the streptomycin/spectinomycin resistance gene of the transformation plasmid backbone were performed to exclude the presence of residual *A*. *tumefaciens* bacteria within the putative transformation lines, interfering with the results of the transformation verification analyses. The T-DNA integration locus was identified via Thermal Asymmetric Interlaced PCR (TAIL-PCR), as described by Liu et al. (1995) and Fladung and Polak (2012). All the PCR primers used are listed in Table S1. The obtained Sanger sequences of the integration loci, provided by StarSeq (Mainz, Germany), were then compared to the two *P*. *tremula* × *P*. *alba* haplotypes (HAP1 and HAP2) using BLAST analysis via the Phytozome database.

### 2.5 Drought stress experimental setup

A three-week greenhouse experiment was conducted from November 22 to December 13, 2023. Rooted head cuttings from the HDG11-2 transformation line and the WT control line were transferred from *in vitro* culture to sieved tree nursery soil. Eight plants from the HDG11-2 line and 14 plants from the WT line were potted to a final pot size of 3 L. The plants were cultivated under controlled conditions with a consistent 16/8 h light/dark regime, 22 °C, and 60% RH. The pots were randomly placed on a greenhouse bench with additional WT plants around the experiment to buffer side effects from environmental fluctuations. When the plants reached a height of approximately 30 cm, drought stress was applied to half of the plants by withholding irrigation, while the other half received a daily regular irrigation of 130 mL, using an automatic drip irrigation system (Rain Bird Corporation; Azusa, CA, USA), which served as the control group for each line. Pot saucers were placed upside down underneath each experimental plant for runoff of excess water and to avoid water accumulation in the root zone. At the beginning of the experiment, all the plants were equally irrigated with 1 L for complete soil saturation. The experimental plants were phenotyped at four sampling days (SD) with SD I = start of the experiment and no drought stress, SD II = 7 days without watering, SD III = 14 days without watering, and SD IV = 7 days of rewatering to target specific stages of the plants’ drought stress and recovery response. Additionally, the soil moisture levels of each pot were monitored on each SD via a micro station data logger and soil moisture sensors (HOBO U21 USB and HOBO EC5 Soil Moisture Smart Sensor; Onset Computer Corp., Bourne, MA, USA). Based on the measured soil volumetric water content (VWC), SD II was defined as moderate stress, corresponding to plants of the stress group grown at approx. 10% VWC, whereas SD III represented severe stress, corresponding to plants of the stress group grown at VWC < 5.

As a second approach, a three-week climate chamber experiment was conducted from January 23 to February 13, 2025, to validate the findings from the earlier greenhouse experiment. Rooted head cuttings from the transformation line HDG11-2 and WT were transferred from *in vitro* culture to soil as previously described. The transferred plants were then acclimatized to the climate chamber conditions (Weiss Technik, Lindenstruth, Germany) in a sealed container with a 16/8 h light/dark cycle with 16 µmol m^−2^ s^−1^ photosynthetic photon flux density (PPFD), 22/17 °C temperature, and 70% RH, with a gradual opening of the lid over a 10-day period. The plantlets were subsequently transferred into 1 L cultivation pots until they reached a height of approximately 30 cm. All plants were then randomly arranged in the climate chamber with unchanged adjustments and approx. 35 µmol m^−2^ s^−1^ PPFD. Here, 10 individuals per line were subjected to drought conditions without irrigation, while six individuals per line received daily manual irrigation with 100 mL of water. The experimental plants were phenotyped, and sampling was carried out at three SD with SD I = start of the experiment and no stress, SD II = 7 days without watering (severe stress), and SD III = 14 days of rewatering (recovery). Due to the smaller pot size in the climate chamber experiment, soil moisture declined more rapidly, resulting in earlier onset of severe drought stress, which was confirmed by VWC measurement during SD II.

### 2.6 Measurement of morphological, physiological, and molecular indicators

#### 2.6.1 Relative water content

To measure the relative water content (RWC), the 3^rd^ fully expanded leaf from each plant was excised in both experiments, specifically under severe stress and rewatering conditions. The cut leaves were immediately weighed to obtain the fresh weight (FW). The leaves were subsequently submerged in a container filled with distilled water to achieve full turgidity. After a period of 24 h, the leaves were removed from the water, excess water was gently blotted off, and the turgid weight (TW) was recorded. The leaves were subsequently dried in an oven at 80 °C for 48 h and then weighed again to obtain the dry weight (DW). The RWC was calculated as RWC = (FW [g] - DW [g]) / (TW [g] - DW [g]) × 100 [%].

#### 2.6.2 Proline quantification

Proline was measured from the 4^th^ fully expanded leaf, according to Bates et al. (1973), and adapted to a microplate-based protocol (Abrahám et al., 2010). In brief, 50 mg of leaf material was homogenized in a bead ruptor under steady cooling. To extract disturbing proteins, each sample was mixed with 1 mL of 5-sulfosalicylic acid and centrifuged at 12,000 × g for 10 min. Two hundred microliters of the supernatant were mixed with ninhydrin, acetic acid, and phosphoric acid and incubated at 96 °C. The reaction was stopped on ice, and each sample was supplemented with 1 mL of toluene to extract the proline-ninhydrin mixture. The determination of proline content was conducted by measuring the absorbance of the toluene proline mixture at 520 nm in a 96-well plate via a microplate reader (SPECTROstar Nano; BMG LABTECH, Ortenberg, Germany). Proline accumulation was determined via the standard curve method.

#### 2.6.3 Malondialdehyde quantification

Lipid peroxidation was determined by quantification of malondialdehyde (MDA) according to Hodges et al. (1999). One hundred milligrams of the 4^th^ fully expanded leaf were homogenized under steady cooling and mixed with 1.5 mL of 0.1% trichloroacetic acid to precipitate proteins. Five hundred microliters were then mixed with reaction solution 1 (comprising 0.01% 2,6-di-ter-butyl-4-methylphenole and 20% trichloroacetic acid) and another 500 µL with reaction solution 2 (comprising reaction solution 1 with additional 0.65% 2-thiobarbituric acid) and subsequently incubated at 96 °C for 30 min. The reaction was stopped on ice and centrifuged at 8,000 × g and 4 °C for 10 min. MDA was determined via photometry at 440, 532, and 600 nm using the microplate reader. The MDA levels were calculated based on the formulas Hodges et al. (1999) and Landi (2017).

#### 2.6.4 Total shoot biomass

To determine the total shoot biomass, all shoots of the plants from the growth chamber experiment were isolated from the below-ground biomass, and the fresh weights of all the leaves and stems per sample were weighed. Afterward, the leaves were oven dried at 80 °C for 48 h, and the stems were oven dried at 105 °C for 24 h, after which the shoot dry weight was measured.

#### 2.6.5 Height and stem diameter

The height of each plant was measured using a standard folding rule from the base of the stem to the tip of the shoot. Additionally, the stem diameter was measured with a vernier caliper at a height of 10 cm above the base of the shoot.

#### 2.6.6 Number of leaves

On each SD, the fully expanded leaves of each individual plant were counted. A fully expanded leaf was defined as a leaf measuring ≥ 2.5 cm in diameter (typically the 2^nd^ or 3^rd^ leaf from the apex). After each SD, the shoot tips above the first fully expanded leaf were labeled to accurately determine the number of newly formed leaves.

#### 2.6.7 Stomatal conductance

Stomatal conductance was measured via an SC-1 leaf porometer (METER Group, Pullman, WA, USA). Measurements were conducted on the 3^rd^ fully expanded leaf of each plant.

#### 2.6.8 Chlorophyll quantification

The chlorophyll content in the leaves was quantified via DUALEX (Force-A, Orsay, France) on the 3^rd^ fully expanded leaf of each plant.

#### 2.6.9 Stomatal density and trichome observation

Trichome removal and stomatal counting were performed according to the protocol described by Wodtke et al. (2025). For each line, three leaves from *in vitro*-grown plantlets were selected for analysis. The stomata were visualized under a light microscope (Axioscope 5; ZEISS, Oberkochen, Germany) and subsequently counted using the ImageJ 1.54p software (Schneider et al., 2012) to determine the stomatal density (number of stomata per mm²). Trichome observation was carried out on *in vitro*-grown plantlets of the two lines using a light microscope (Stemi 2000-C; ZEISS, Oberkochen, Germany).

### 2.7 Gene expression analyses

Quantitative real-time PCR (RT-qPCR) was performed to analyze the expression levels of selected genes, based on their functional relevance to the observed phenotypic responses, including *PtaHDG11*, *GLABRA 2* (*GL2*), and *STOMATAL DENSITY AND DISTRIBUTION 1* (*SDD1*), as well as *A*. *thaliana* homologs of *SUPEROXIDE DISMUTASE 2* (*SOD2-like*), *CATALASE 2* (*CAT2-like*), *XYLOGLUCAN:XYLOGLUCOSYL TRANSFERASE 32* (*XTH32-like*), and *α-EXPANSIN-15* (*EXPA15-like*) identified through BLAST analysis in *P. tremula* × *P. alba* and designated as ‘-like’ genes, in the WT and the transgenic line. Total RNA was isolated from the leaf tissue harvested from three biological replicates per line during the severe stress (SD III) in the greenhouse experiment, with sampling performed in the morning, at approx. 10 a.m. The RNA isolation was performed according to Bruegmann et al. (2024), using the innuPREP Plant RNA Kit (IST Innuscreen; Berlin, Germany). cDNA synthesis and RT-qPCR were carried out via the GoTaq 2-Step RT-qPCR System (Promega Corp.; Madison, WI, USA) following the manufacturer’s instructions. The C1000 Touch Thermal Cycler (Bio-Rad Laboratories Inc.; Hercules, CA, USA) was programmed with the following settings: initial denaturation at 95 °C for 2 min, followed by 40 cycles of 95 °C for 15 s and 58 °C for 1 min. The specific amplification of each target gene was analyzed using a melting curve (95 °C for 15 s, 59 °C for 1 min). The relative expression of each target gene was calculated based on the ΔΔCt-method (Livak and Schmittgen, 2001). All measurements were normalized to the poplar UBIQUITIN (*PtaUBQ*) housekeeping gene (Regier and Frey, 2010). ΔΔCt values were calculated relative to the calibrator defined as the mean ΔCt of the WT control group. To ensure data quality, a leave-one-out outlier rule was applied per genotype × treatment group where a value was removed if its absolute deviation from the group median exceeded 0.75 cycles and was at least twice the second-largest deviation within that group. Statistical analysis was performed on ΔΔCt values using a linear model (ΔΔCt ∼ Genotype × Treatment). Estimated marginal means were back-transformed to the fold-change scale for reporting and visualization.

### 2.8 Statistical analyses

All analyses were performed in R version 4.4.2 and RStudio version 2024.12.0.467. Repeated-measure traits were modeled using linear mixed-effects models (lme4, lmerTest) with line, treatment, and SD as fixed effects and plant individual as a random intercept. For each trait, model fit and residual diagnostics (Shapiro–Wilk, Levene’s test, DHARMa) were evaluated for both raw and log+1-transformed data. The transformation yielding residuals closest to normality and homoscedasticity was selected for final modeling. Fixed effects were assessed by Type III ANOVA, with pairwise contrasts of estimated marginal means adjusted by Holm’s method. Traits measured only once were analyzed using univariate tests (Student’s t-test or Welch’s t-test), depending on homogeneity of variances. Outliers were removed within each subline and treatment group for each trait using the interquartile range (IQR) method. Statistical significance was set at α = 0.05. Pearson’s correlation coefficient (r) was used to assess linear relationships between physiological, morphological, and biochemical traits, whereas Spearman’s rank correlation coefficient (ρ) was used to assess monotonic relationships.

## 3 Results

The BLAST search of the *At*HDG11 protein sequence identified two paralogous genes in both alleles of *P. tremula* × *P. alba* (Fig. 1A). The analysis revealed that the gene *PtXaAlbH.15G026700* from the hybrid’s *P. alba* allele exhibited the highest protein similarity (70.9%). The Maximum Likelihood phylogenetic analysis of the HD-ZIP IV subfamily revealed that the *AtHDG11*/*HDG12* cluster and the corresponding *Populus* cluster diverged from a common ancestor with high statistical support (Bootstrap 99), confirming the orthologous relationship of the entire *Populus* group consisting of both *P*. *tremula* × *P. alba* alleles and *P*. *trichocarpa* (Fig. 1B). *PtXaAlbH.15G026700,* the closest homolog, was selected as the candidate gene, referred to as *PtaHDG11*.

**Fig. 1.**
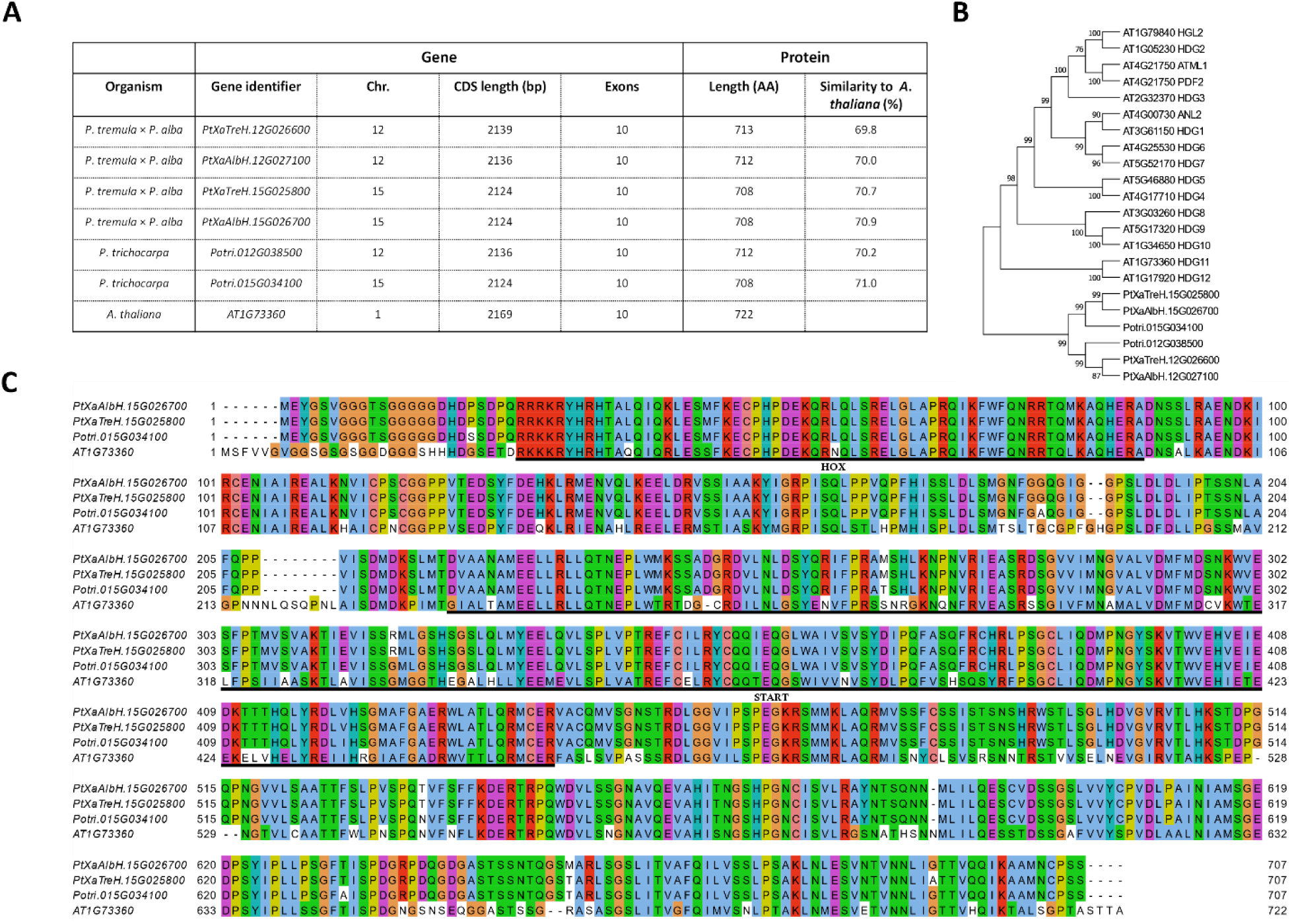
Identification of the candidate gene *PtaHDG11*. (A) Overview of genetic and protein-related data of *AtHDG11* homologs identified via BLAST analysis, including protein similarity. (B) Phylogenetic tree with all *A*. *thaliana* HD-ZIP IV genes and all identified poplar homologs of *AtHDG11.* (C) Multiple protein sequence alignment (CLUSTALW) of the *HDG11* gene from both *P. tremula* × *P. alba* alleles (*PtXaAlbH.15G026700*, *PtXaTreH.15G025800*), *P. trichocarpa* (*Potri.015G034100*), and *A. thaliana* (*AT1G73360*), visualized in Jalview using ClustalX color scheme. Colors indicate AA physiochemical classes (blue: hydrophobic, green: polar, red: positively charged, purple: negatively charged, pink: cysteine, brown: glycine, yellow: proline, turquoise: aromatic), highlighting conserved residues and conservative substitutions. The two HD-ZIP IV protein domains HOX and START are underlined in black

A comparative analysis of the protein sequences of *HDG11* genes was conducted via multiple sequence alignment, incorporating the HDG11 protein sequences of *P. tremula* × *P. alba*, *A. thaliana*, and *P. trichocarpa* (Fig. 1C). The two *P. tremula* × *P. alba* alleles presented only 3 amino acid (AA) differences within the 707 AA protein, indicating a high degree of similarity between them. The comparison between the species revealed several AA differences between the groups. Specifically, 68% of the AA sequence of the poplar hybrid was identical to that of *A. thaliana*, with 256 AA differences observed between the two species. The alignments of the *P. tremula* × *P. alba* alleles and *P. trichocarpa* alleles were more similar, with differences in 8 AAs between *P. trichocarpa* and the *P. tremula* allele and in 9 AAs compared with the *P. alba* allele. Differences occurred mainly at the N- and C-termini or outside of conserved protein domains. Notably, the protein sequences of the two poplar species presented nearly perfect similarity within the conserved protein domains of HOX and START, which classify the HD-ZIP IV class, with only two exceptions in the START domain at AA positions 264 and 320. In total, the sequence identity between *P. tremula* × *P. alba* and *A*. *thaliana* reached approx. 95% in the HOX and approx. 89% in the START domain. Interestingly, the AA differences in the HOX domain between the poplar species and *A*. *thaliana* occurred mostly in the first half of the domain, whereas the second half, which includes the helix III region (Nakamura et al., 2006), was identical across all species, except for an AA change from leucine in *A. thaliana* to methionine in poplars.

*Agrobacterium*-mediated transformation and subsequent *in vitro* regeneration yielded three individual lines, which were confirmed to be transgenic by PCR analysis of the *nptIIo* marker gene and the transgenic *PtaHDG11*. However, subsequent Southern blot analysis revealed double T-DNA integration only in HDG11-2. In contrast, HDG11-1 and HDG11-3 could not be demonstrated to carry stable T-DNA insertions. Therefore, only line HDG11-2 was considered for all the following experiments. Finally, the T-DNA integration events detected in HDG11-2 via TAIL-PCR occurred exclusively in noncoding regions of the genome.

### 3.1 Greenhouse stress experiment

The WT and HDG11-2 lines were subsequently used for a greenhouse drought stress experiment to assess their performance under water-limited conditions. The greenhouse drought stress experiment was conducted for 21 days, with a SD every 7^th^ day. The measurement of soil moisture revealed no significant differences in VWC between the two lines within a treatment group on any SD and was around 10% VWC on SD II and < 5% on SD III for both lines within the stress treatment as well as above 20% throughout the whole experiment for both lines in the control treatment (Fig. S2).

At SD II, under moderate stress, both lines of the stress treatment began to exhibit signs of stress, but significant growth cessation was not observed until the severe stress phase on SD III. In the control groups, a steady increase in plant height of approximately 30 cm was observed throughout the experiment (Fig. 2A). In contrast, the stress treatments led to a significant reduction in plant height compared with that of the corresponding control groups on the severe stress and recovery SDs, with the WT showing the greatest difference (HDG11-2 p < 0.01; WT p < 0.001). Although no significant differences in plant height or stem diameter were found between the tested lines within a treatment on a given SD, the stress treatments resulted in a consistent pattern of growth reduction in both lines. On SD IV, under recovery conditions, HDG11-2 presented a mean increase in height of approximately 4 cm greater than that of the WT. The analysis of stem diameter revealed significant differences between the same line under different treatments. Under severe stress, compared with the control plants, the WT and HDG11-2 plants presented reduced stem diameters (p < 0.05, Fig. 2B). For the HDG11-2 line, this effect was consistent on SD IV (p < 0.05).

**Fig. 2.**
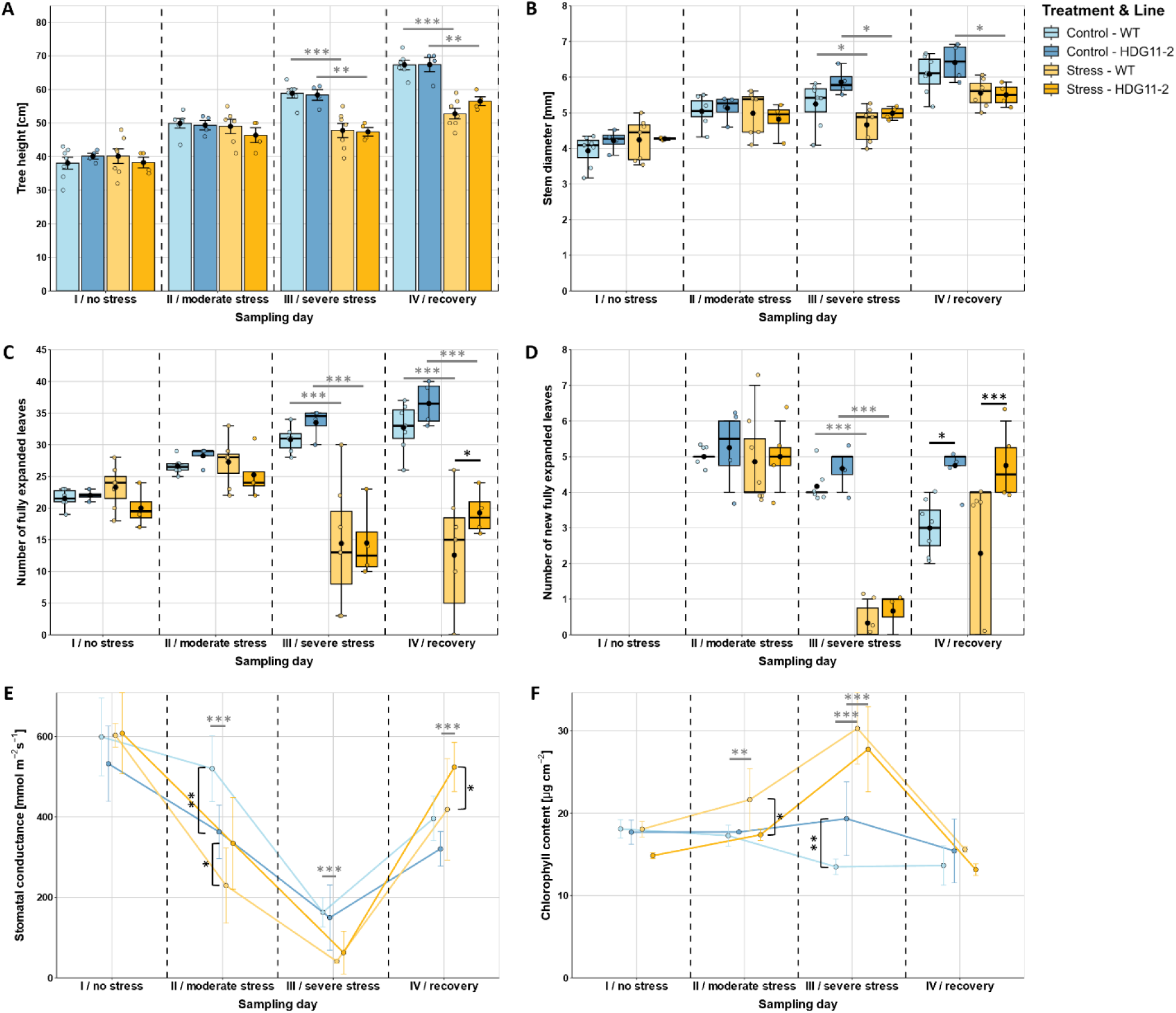
Morphological and physiological responses of WT and HDG11-2 plants under drought stress conditions in the greenhouse. Measurements of (A) tree height, (B) stem diameter, (C) number of fully expanded leaves, (D) number of new fully expanded leaves, (E) stomatal conductance, and (F) chlorophyll content. The graphs present the means ± standard errors (SEs; n_WT_ = 7, n_HDG11-2_ = 4). Black asterisks indicate significant differences between lines within the same treatment, whereas grey asterisks indicate significant differences between treatments within the same line for each SD, based on linear mixed-effects models with Holm-adjusted emmeans post hoc contrasts (* p < 0.05, ** p < 0.01, *** p < 0.001). Comparisons among SD are reported in the main text

Similar to tree height and stem diameter, the number of fully expanded leaves increased steadily in the control groups of both lines (Fig. 2C). Compared with their respective control plants, the stress-treated plants presented a significant reduction in leaf number under severe stress and recovery conditions (p < 0.001). However, during recovery on SD IV, former stressed HDG11-2 presented a significantly greater leaf number than did the former stressed WT (19 vs. 12; p < 0.05).

Additionally, we counted the number of new fully expanded leaves, which represents the potential for leaf growth under or after drought stress conditions (Fig. 2D). Under severe stress on SD III, both the WT and HDG11-2 lines presented severe reductions in new leaf formation compared with their respective controls (p < 0.001). During the 7-day recovery period, three out of seven former stress-treated WT trees failed to produce new leaves. In contrast, HDG11-2 recovered its ability to form new leaves in all individuals, reaching nearly the levels observed during moderate stress. Here, HDG11-2 outgrew the WT by 3 new fully expanded leaves (p < 0.001). Interestingly, the WT control plants presented a reduced number of new fully expanded leaves over the course of the experiment, resulting in a significant difference from the control plants of HDG11-2 at SD IV (p < 0.05) and also a significant difference from their own new fully expanded leaves at SD II (p < 0.01).

The stomatal conductance was measured on each SD on the 3^rd^ fully expanded leaf of each individual (Fig. 2E). Both lines in the stress treatment presented a significant reduction in stomatal conductance during moderate stress on SD II (p < 0.001 for both lines) and severe stress on SD III (p < 0.001 for both lines). On both SDs, the stomatal conductance of the stressed WT significantly differed from those of its own control group (p < 0.001 on both SDs), whereas no such significant differences were observed for the HDG11-2 line. During SD II, stressed WT plants presented stomatal conductance values ranging from 96 to 366 mmol m^−2^ s^−1^, which significantly differed from those of HDG11-2, which ranged from 204–480 mmol m^−2^ s^−1^ (p < 0.05). During SD III, the stomatal conductance further decreased to a range of values between 36–50 mmol m^−2^ s^−1^ in the WT and 23–140 mmol m^−2^ s^−1^ in HDG11-2, with the transgenic line displaying a broader range of stomatal conductance than the WT. Interestingly, although the control plants of both lines were steadily watered, a reduction in leaf stomatal conductance could also be observed in the same trend as the stressed plants, although not as intense. In the control treatment, a significant difference between WT and HDG11-2 was observed at SD II (p < 0.01). Upon recovery, the previously stressed HDG11-2 plants presented significantly greater stomatal conductance than the control plants did (p < 0.001), indicative of a compensatory response that surpassed the stress-free state of the plants. Here, the stomatal conductance values ranged between 456-604 m^−2^ s^−1^, that also led to a significantly difference to the previously stressed WT plants, where values ranged between 213-550 m^−2^ s^−1^ (p < 0.05).

The chlorophyll content was measured on each SD in the 3^rd^ fully expanded leaf of all individuals (Fig. 2F). Compared with its own control group, the chlorophyll content in the WT line significantly increased in the stress treatment group on SD II (p < 0.01) and on SD III for both lines (p < 0.001). The elevated chlorophyll content of the stress treated WT line on SD II also resulted in a significant difference compared to the stress treated HDG11-2 line (p < 0.05). Further, the control plants presented significant differences, where the WT plants showed a significant decrease in chlorophyll content during SD III compared to the HDG11-2 line (p < 0.01).

As evident from the data presented in Fig. 2A–F, the drought stress treatment had a pronounced effect on both lines of the stress group. Additional phenotypic examinations revealed that during recovery, some of the former stressed WT plants exhibited irreversible shoot tip damage and complete leaf loss, while all HDG11-2 plants fully regained turgor and vitality, demonstrating their superior drought resilience.

The mean RWC of the 3^rd^ fully expanded leaf remained consistently above 80% for both control lines throughout the experiment (Fig. 3A). A significant difference in RWC was observed between the WT and HDG11-2 lines in the stress group during SD III (p < 0.01). In general, the RWC of both stressed lines decreased under severe stress, with 68% in the WT and 81% in the HDG11-2 line, resulting in significant differences in comparison to their own control groups (p < 0.001; p < 0.05, respectively). During recovery, the RWC increased again in both lines, where the significant difference in RWC between the WT and HDG11-2 plants in the stress treatment group was still consistent (p < 0.01). Here, HDG11-2 reached a level of 96%, surpassing the RWC level of the own control group (88%).

**Fig. 3.**
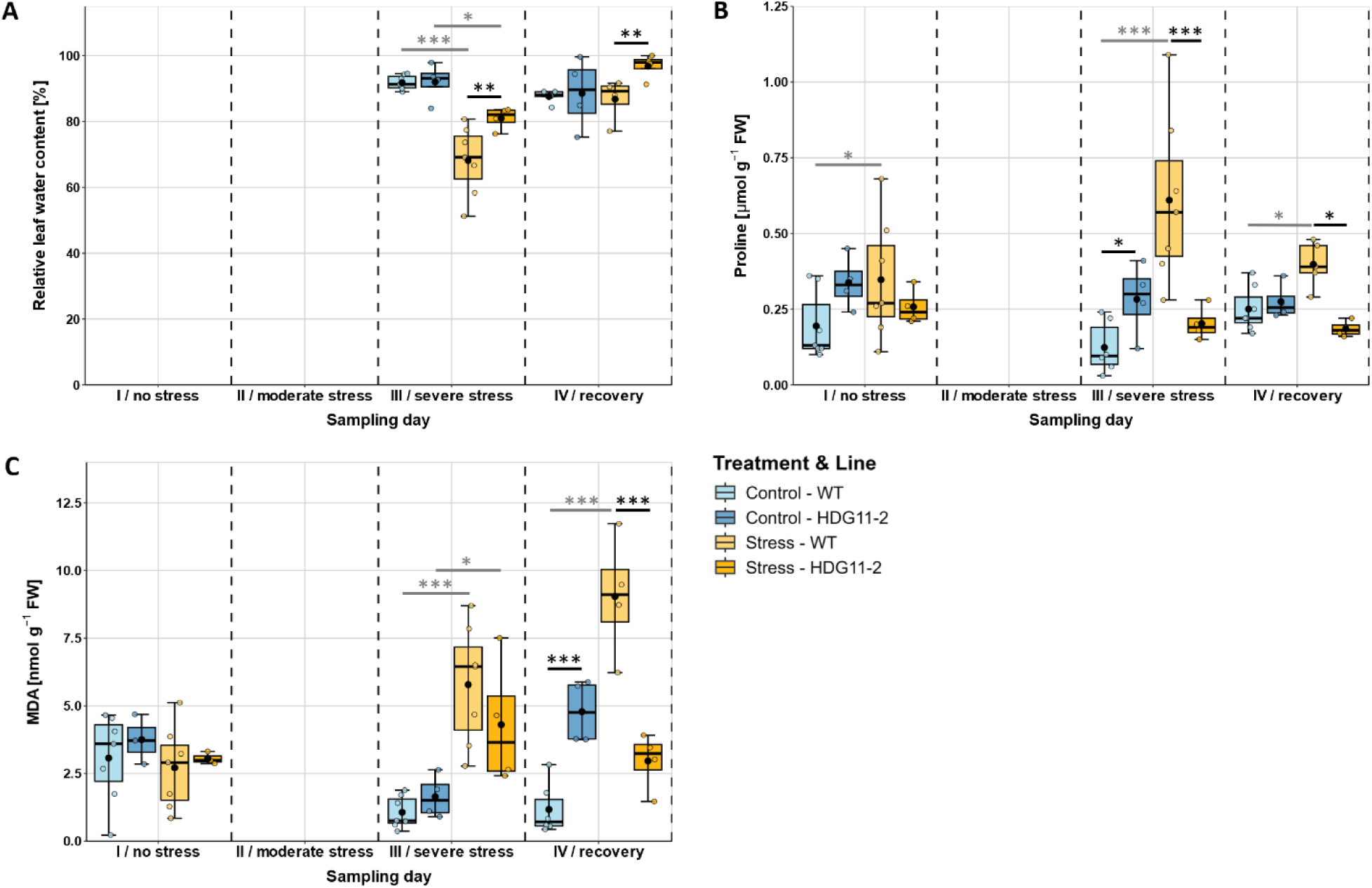
Molecular physiological responses of WT and HDG11-2 plants under drought stress conditions in the greenhouse. Measurement of (A) RWC, (B) proline content, and (C) MDA content. The graphs present the means ± SEs (n_WT_ = 7, n_HDG11-2_ = 4). Black asterisks indicate significant differences between lines within the same treatment, whereas grey asterisks indicate significant differences between treatments within the same line for each SD, based on linear mixed-effects models with Holm-adjusted emmeans post hoc contrasts (* p < 0.05, ** p < 0.01, *** p < 0.001). Comparisons among SD are reported in the main text

Changes at the molecular level were observed by measuring the accumulation of the stress-triggered molecules proline and MDA during the experiment under the two treatments and in the two different lines. Proline and MDA contents were measured on SD I (no stress) as well as during severe stress and recovery (SD II and SD III). The basal mean proline concentration ranged from 0.1 to 0.35 µmol g^−1^ FW for both lines in both treatments at the beginning of the experiment (Fig. 3B). These concentrations remained relatively stable throughout the experiment, with the exception of the WT line in the stress treatment. During severe stress on SD III, proline concentrations increased significantly in the WT line, reaching a mean concentration of 0.61 µmol g^−1^ FW, which was highly significantly different from that in the HDG11-2 overexpression line and the own control group of the WT (both with p < 0.001). Although the proline concentration of the WT decreased during recovery on SD IV, the differences were still significant compared to HDG11-2 and the WT control group (p < 0.05; p < 0.05, respectively). Initially, both lines presented low mean MDA accumulations (< 5 nmol g^−1^ FW) in both treatment groups, which remained stable in the control groups throughout the experiment (Fig. 3C). Under severe stress on SD III, the MDA concentrations increased slightly in both stressed lines, with significant differences compared with their respective controls (HDG11-2 p < 0.05; WT p < 0.001). Notably, the stress group of the WT line further presented a strong increase in MDA concentration during recovery on SD IV, with a mean value of 9.04 nmol g^−1^ FW, which was highly significantly different from that of the overexpression line HDG11-2 (2.97 nmol g^−1^ FW, p < 0.001) and its own control group (1.18 nmol g^−1^ FW, p < 0.001).

### 3.2 Trichome phenotyping

During the first experiment in the greenhouse, we observed that the individuals of the HDG11-2 line presented a glabrous phenotype characterized by a lack of trichomes on both the abaxial leaf side and the stem. This phenotype was also observed in *in vitro*-grown plantlets of the HDG11-2 line. Light microscopic examination confirmed the complete loss of trichomes in these individuals, whereas WT individuals presented typical trichome density (Fig. 4).

**Fig. 4.**
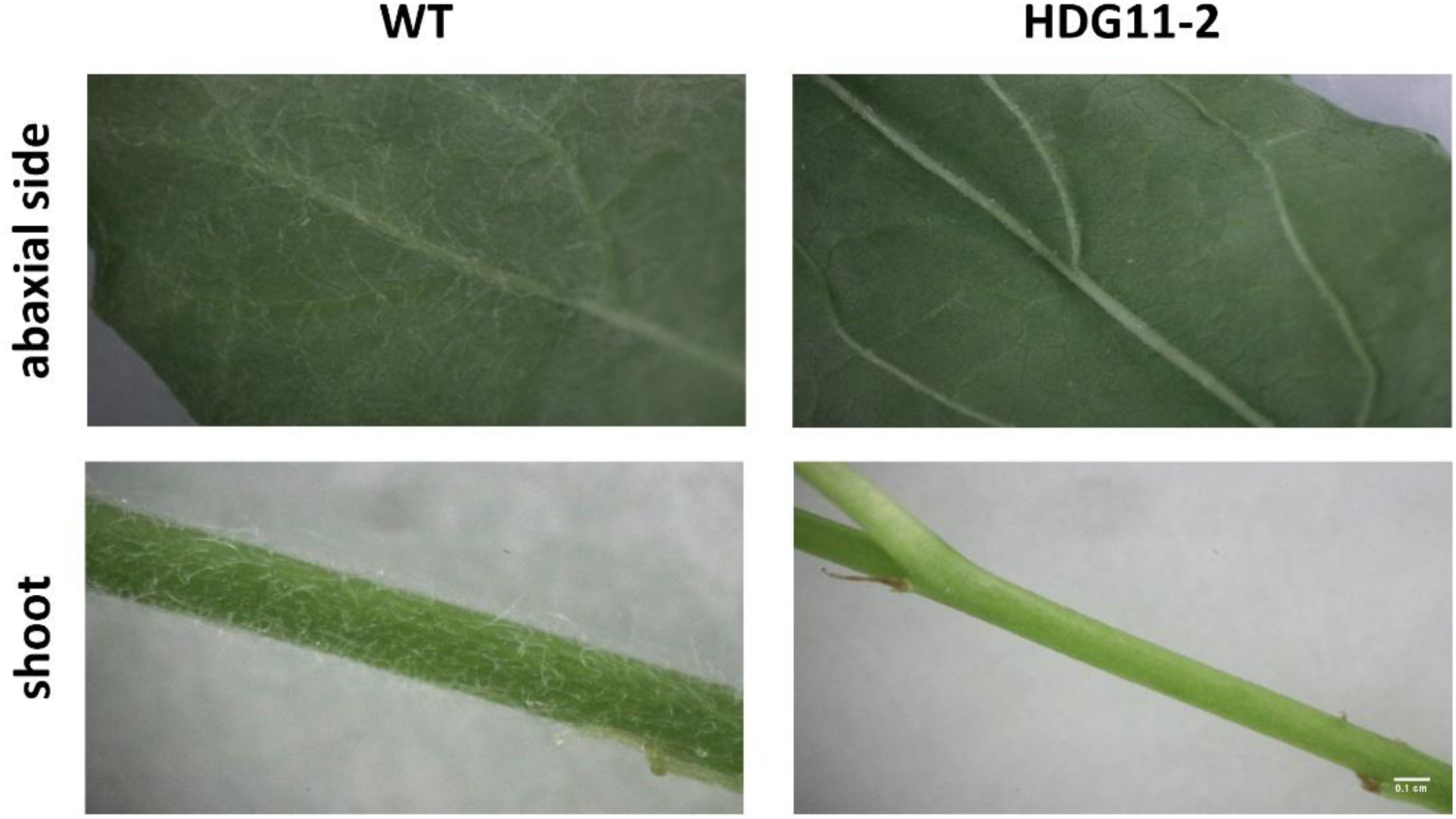
Microscopic examination of the abaxial leaf surface and stem of *in vitro*-grown WT and HDG11-2 plants revealed a distinct glabrous phenotype in HDG11-2. Trichomes were absent in all examined HDG11-2 plants but consistently present in all WT plants. Because the phenotype was uniform within both groups, no statistical analysis was performed

#### Gene expression analyses

To better contextualize the effects of increased drought-stress tolerance and the absence of trichomes in line HDG11-2, we performed RT-qPCR analyses. The expression of the candidate gene *PtaHDG11* and of several target genes was analyzed, including *PtaSOD2*-*like* and *PtaCAT2*-*like*, which are associated with antioxidant defense, *PtaSDD1*, which is involved in stomatal density and distribution, *PtaGL2*, which is associated with trichome development, and *PtaXTH32*-*like* and *PtaEXPA15*-*like*, which are involved in cell wall loosening and remodeling. Expression of *PtaHDG11*, expressed as fold change relative to the WT control, was significantly higher in the transgenic line HDG11-2 than in WT under severe stress on SD III (p < 0.05; Fig. 5A). Under control conditions, *PtaHDG11* expression was also approximately 1.6-fold higher in HDG11-2 than in WT, although this difference was not significant. While the expression decreased under severe stress in both lines, only WT showed significant downregulation relative to its control (p < 0.05).

**Fig. 5.**
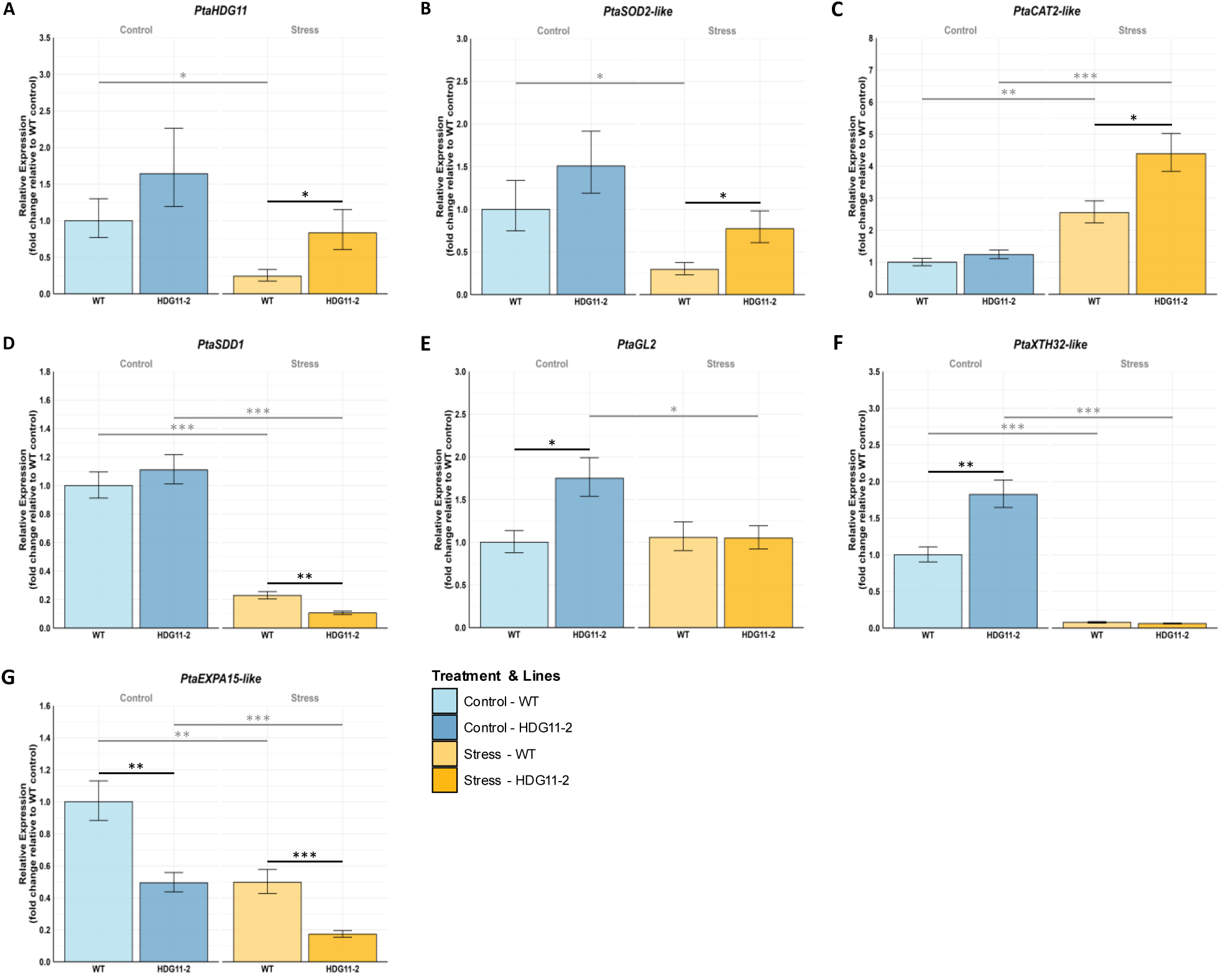
Gene expression analyses of WT and HDG11-2 on SD III in the greenhouse experiment. Expression analyses of (A) *PtaHDG11*, (B) *PtaSOD2-like*, (C) *PtaCAT2-like*, (D) *PtaSDD1*, (E) *PtaGL2*, (F) *PtaXTH32-like*, and (G) *PtaEXPA15-like*. Bars show model-based mean relative expression levels (fold change relative to the WT control calibrator) ± SE (n = 3). Black asterisks indicate significant differences between lines within the same treatment, whereas grey asterisks indicate significant differences between treatments within the same line, based on contrasts of estimated marginal means from a linear model fitted to ΔΔCt values (ΔΔCt ∼ Genotype × Treatment) with * p < 0.05,** p < 0.01, and *** p < 0.001

Given the enhanced drought stress tolerance of HDG11-2 in the greenhouse experiment, we next examined the expression of antioxidant genes. Similar to *PtaHDG11*, expression of *PtaSOD2*-*like* was approximately 1.5-fold higher in HDG11-2 than in WT under severe stress (p < 0.05; Fig. 5B), whereas only WT showed a significant reduction relative to its control (p < 0.05). For *PtaCAT2*-*like*, both WT and HDG11-2 showed significantly increased expression under severe stress compared with their respective controls (WT: p < 0.01; HDG11-2: p < 0.001). Under severe stress, *PtaCAT2*-*like* expression was also significantly higher in HDG11-2 than in WT, corresponding to an approximately 1.8-fold increase (p < 0.05; Fig. 5C). Overall, the expression patterns of the antioxidant genes *PtaSOD2*-*like* and *PtaCAT2*-*like* suggest that transcript levels were maintained more strongly in HDG11-2 than in WT under stress conditions, with a tendency toward higher *PtaSOD2*-*like* expression already under control conditions.

Expression of *PtaSDD1* was highly significantly downregulated under severe stress in both lines (p < 0.001; Fig. 5D). In addition, HDG11-2 showed significantly lower *PtaSDD1* expression than WT under severe stress (p < 0.01). Because *PtaSDD1* is involved in the regulation of stomatal density, its strong downregulation in HDG11-2 might suggest an increase in stomatal density. However, subsequent microscopic analysis of in vitro-grown WT and HDG11-2 plants revealed no significant difference in stomatal density between the two lines (Fig. S3).

Expression analysis of *PtaGL2*, which is associated with trichome development, revealed significantly higher expression in HDG11-2 than in WT under control conditions, suggesting altered regulation of trichome formation (Fig. 5E). However, *PtaGL2* expression was significantly reduced under stress and reached nearly the same level in HDG11-2 as in WT, where the *PtaGL2* expression was not affected by the stress treatment (p < 0.05). In addition, *PtaXTH32*-*like*, a xyloglucan endotransglycosylase/hydrolase involved in cell wall plasticity and extensibility, was significantly higher expressed in HDG11-2 than in WT under control conditions (p < 0.01), whereas expression decreased drastically under severe stress in both lines (p < 0.001; Fig. 5F).

In contrast, expression of *PtaEXPA15*-*like*, an α-expansin gene involved in cell wall loosening, was significantly lower in HDG11-2 than in WT under both control and severe stress conditions (p < 0.01 and p < 0.001, respectively; Fig. 5G). In addition, *PtaEXPA15*-*like* expression decreased significantly under stress in both WT and HDG11-2 relative to their respective controls (WT: p < 0.01; HDG11-2: p < 0.001).

### 3.3 Climate chamber stress experiment

To further test and validate the increased drought stress tolerance observed in the greenhouse experiment under a different environment, a climate chamber drought stress experiment was conducted on lines HDG11-2 and WT, with a higher number of individuals in the stress treatment (n = 10) for both lines. Phenotypic examinations of the former stress treated plants at the end of the recovery phase (SD III) revealed distinct differences in the drought tolerance/recovery ability of both lines (Fig. 6A). The WT line exhibited pronounced leaf loss and partially non-recoverable shoot tip damage, whereas the HDG11-2 line remained largely vital. Similar to the first experiment, no significant differences in tree height were observed between WT and HDG11-2 under the same treatment conditions (Fig. 6B). The height of the control plants in both lines steadily increased, with an average increase of approximately 12 cm over the course of the experiment. In contrast, during severe stress on SD II, plants of both lines subjected to drought stress presented cessation of growth, resulting in significant differences compared with their respective control lines (WT p < 0.05; HDG11-2 p < 0.01), which persisted during the recovery phase on SD III (p < 0.01). However, both lines in the stress group resumed growth during the recovery period, ultimately achieving a mean height increase of approximately 10 cm. Similar results were observed for stem diameter, although significant differences were noted between HDG11-2 and the stress group of WT during recovery, with HDG11-2 exhibiting a thicker stem (p < 0.05, Fig. 6C). Overall, the HDG11-2 line initially presented a lower stem diameter and tree height than the WT line did (albeit not significantly), ultimately surpassing the WT line in terms of these growth parameters by the end of the experiment.

**Fig. 6.**
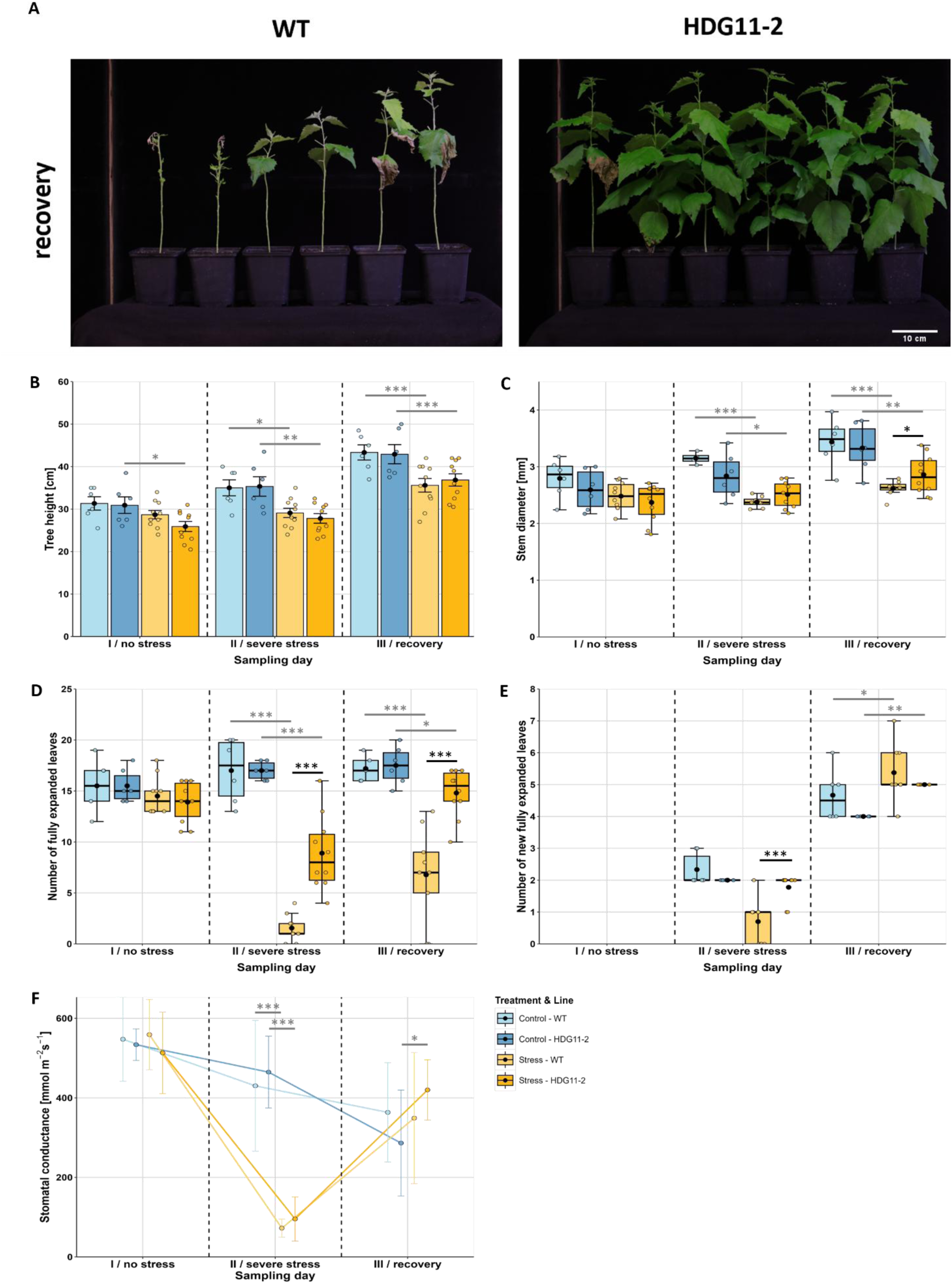
Morphological and physiological responses of WT and HDG11-2 plants under drought stress conditions in the climate chamber. Measurements of (A) Mean six individuals of WT and HDG11-2 after 14 days of rewatering (SD III, recovery) following the stress period, (B) tree height, (C) stem diameter, (D) number of fully expanded leaves, (E) number of new fully expanded leaves, and (F) stomatal conductance. The graphs present the means ± SEs (control treatment, n = 6; stress treatment, n = 10). Black asterisks indicate significant differences between lines within the same treatment, whereas grey asterisks indicate significant differences between treatments within the same line for each SD, based on linear mixed-effects models with Holm-adjusted emmeans post hoc contrasts (* p < 0.05, ** p < 0.01, *** p < 0.001). Comparisons among SD are reported in the main text

A pronounced phenotypic difference emerged between WT and HDG11-2 within the stress treatment group (Fig. 6D). The number of fully expanded leaves differed highly significant between the two lines during severe stress on SD II and the following recovery phase on SD III. On average, HDG11-2 presented 7 more leaves per individual (p < 0.001). Significant differences in the number of fully expanded leaves within the same line and between the two treatments were strongly apparent on SD II for both lines (p < 0.001). During recovery on SD III, this effect persisted in WT, whereas the previously stressed HDG11-2 plants showed a reduced difference relative to its control group (p < 0.05), underscoring the robust recovery of growth parameters in the HDG11-2 overexpression line. This result is further supported by the examination of newly developed leaves (Fig. 6E). Compared with WT plants, HDG11-2 plants presented significantly more new leaves in the stress group during severe stress on SD II (p < 0.001). Additionally, the number of new fully expanded leaves in HDG11-2 did not differ between the two treatments during severe stress, in contrast to the WT (p < 0.001). On SD III, during recovery, both former stressed lines surpassed their respective control (WT: p < 0.05, HDG11-2: p < 0.01).

The stomatal conductance followed the trends observed during the greenhouse experiment. Severe drought stress led to a reduction in stomatal conductance in both stressed lines, with mean values of 72 mmol m^−2^ s^−1^ in WT and 95 mmol m^−2^ s^−1^ in HDG11-2 (Fig. 6F, p < 0.001). During the recovery phase, stomatal conductance increased again in both former stressed lines, with HDG11-2 exhibiting a significant increase that surpassed that of the control group (p < 0.05). Although less pronounced than that in the greenhouse experiment, the stomatal conductance also decreased in the control treatment lines throughout the experiment.

To gain insight into the water content and available water in the plants’ leaves, the mean RWC was measured in the 3^rd^ fully expanded leaf of each individual (Fig. 7A). The RWC decreased in both lines of the stress treatment on SD II, where the WT showed a stronger decrease compared to its control group than HDG11-2 (p < 0.001; p < 0.05, respectively). Also, a significant difference in RWC was observed between the WT and HDG11-2 lines within the stress treatment. Compared with the WT line, the overexpression line presented, on average, a 17% greater RWC (p < 0.001). Additionally, VWC measurements of the soil moisture revealed a significantly higher VWC in the stressed HDG11-2 line (Fig. S4).

**Fig. 7.**
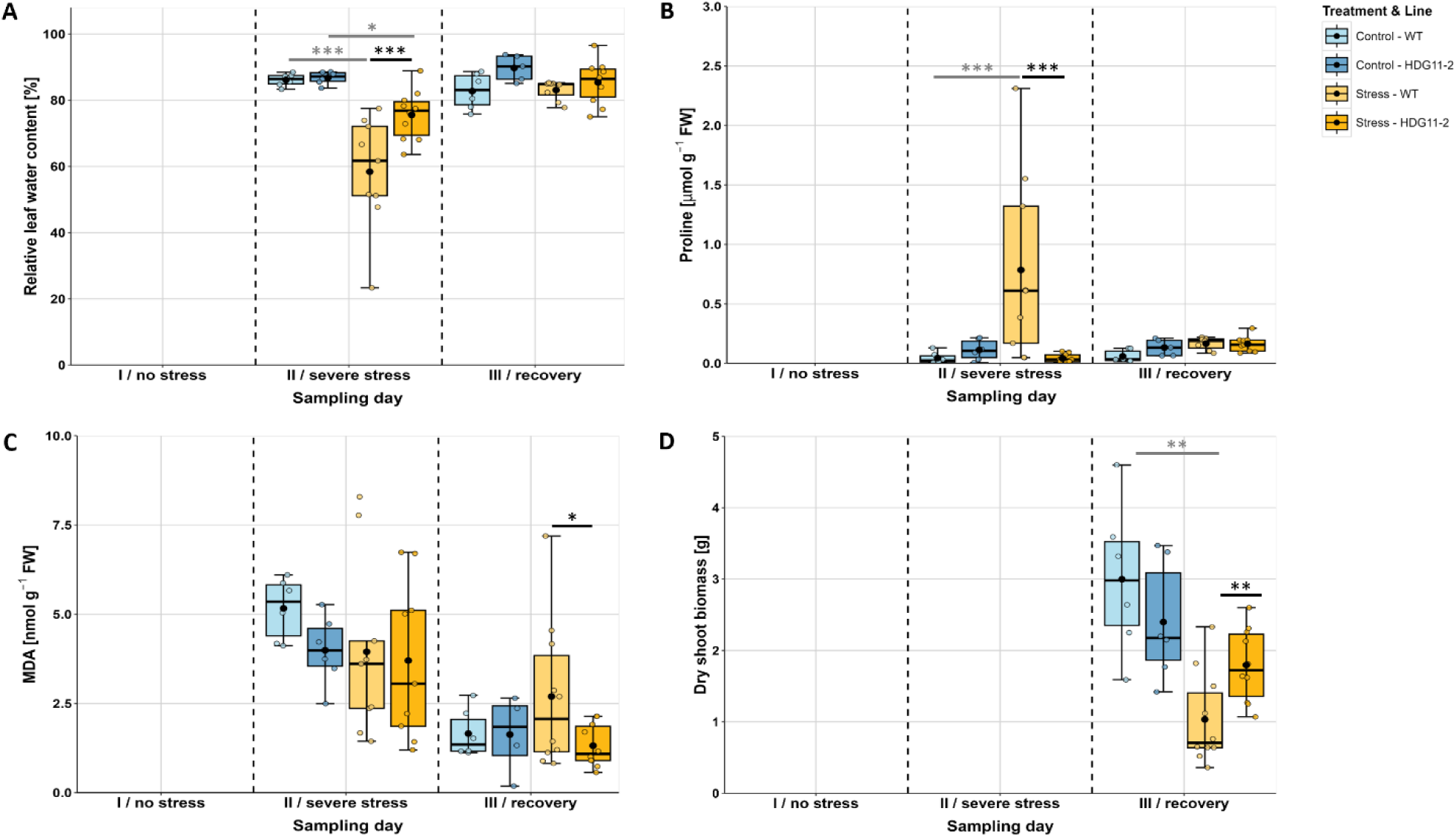
Molecular physiological and physiological responses of WT and HDG11-2 plants under drought stress conditions inside the climate chamber. Measurements of (A) RWC, (B) proline content, (C) MDA content, and (D) dry shoot biomass. The graphs present the means ± SEs (control treatment, n = 6; stress treatment, n = 10). Black asterisks indicate significant differences between lines within the same treatment, whereas grey asterisks indicate significant differences between treatments within the same line for each SD, based on linear mixed-effects models with Holm-adjusted emmeans post hoc contrasts (* p < 0.05, ** p < 0.01, *** p < 0.001). Comparisons among SD are reported in the main text

The mean proline concentration was low, around 0.11 µmol g^−1^ FW, for both lines and treatments on all the SDs, except for the stress-treated WT on SD II (Fig. 7B). Here, the proline concentration increased to a mean value of 0.78 µmol g^−1^ FW, resulting in a highly significant difference compared with that of the stressed HDG11-2 line and its own control group (p < 0.001). Notably, the proline concentration decreased during the recovery phase, returning to a relatively low level. In contrast, a significant difference in the MDA concentration was observed between the WT line and HDG11-2 on SD IV, where the former stressed WT could not decrease the MDA concentration as fast as HDG11-2 (p < 0.05, Fig. 7C).

After 14 days of recovery, the aboveground biomass of all the plants was harvested, oven-dried, and weighed (Fig. 7D). The results revealed that the HDG11-2 overexpression line presented significantly greater dry shoot biomass (p < 0.01), which was consistent with the morphological parameters measured earlier. These findings further support the enhanced drought stress tolerance of the HDG11 overexpression line.

A significant positive linear correlation was observed between the RWC and VWC under severe drought stress on SD II in the HDG11-2 line of the stress group (Pearson’s r = 0.93, p < 0.001; Fig. 8A). Plants with higher soil moisture contents consistently showed higher internal water contents. In contrast, the RWC and VWC of the WT line were not correlated (Pearson’s r = −0.022, p = 0.955). The data revealed distinct clustering by genotype, with the WT plants showing a distribution over the lower range of VWC values (spanning from < 1% to 7%) and the HDG11-2 line primarily distributed within a higher VWC range (spanning almost exclusively from 3% to 15%). Furthermore, at comparable VWC levels, the HDG11-2 line generally maintained a higher RWC than did the WT.

**Fig. 8.**
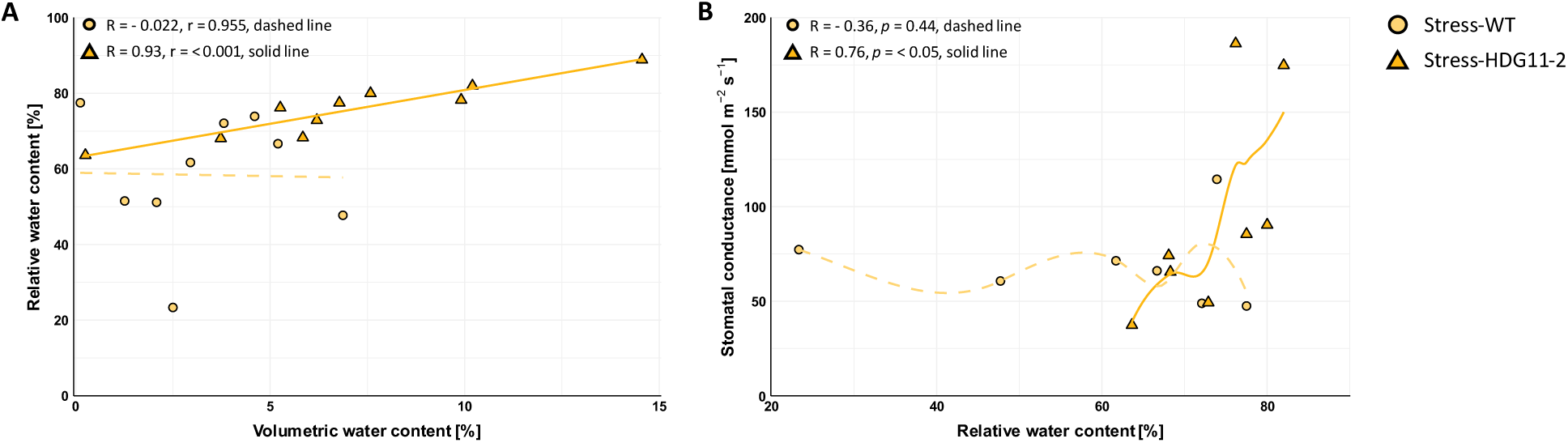
Correlations between the WT and HDG11-2 lines under severe stress in the climate chamber experiment. (A) Pearson correlation: VWC and RWC were significantly positively correlated in the HDG11-2 line, indicating a more stable water status. (B) Spearman correlation: RWC and stomatal conductance showed a significant positive correlation in the HDG11-2 line, suggesting coordinated regulation of water balance and gas exchange in HDG11-2. Significant differences between lines are based on Student’s t-test. n = 7–10

Additionally, a significant positive correlation was detected between RWC and stomatal conductance under severe drought stress in the HDG11-2 line (Fig. 8B). The HDG11-2 line presented a positive correlation with Spearman’s ρ = 0.76 (p < 0.05), maintaining greater RWC and stomatal conductance than those of the WT under the same stress conditions. The data points for the HDG11-2 line were clustered in the right quadrants of the plot, indicating a steeper relationship between RWC and stomatal conductance, whereas the WT data points were located almost horizontally over the whole RWC range with a stomatal conductance almost exclusively below 80 m^−2^ s^−1^.

## 4 Discussion

The poplar-specific *PtaHDG11* overexpression was evaluated in two independent drought stress experiments, utilizing similar experimental designs across distinct environments. These complementary experiments served to assess the reproducibility of the observed responses and to extend the current understanding of the HDG11 TF, particularly by contrasting these findings with the divergent outcomes previously reported for heterologous *AtHDG11* overexpression in *P*. × *tomentosa*. Here, we demonstrated the increased drought stress tolerance in a transgenic *P*. *tremula* × *P*. *alba* line constitutively overexpressing the endogenous *PtaHDG11* gene. Comprehensive phenotyping, which encompasses morphological, physiological, molecular physiological, and genetic aspects, revealed several parameters associated with improved drought stress tolerance in the genetically modified poplars.

### 4.1 Growth and morphological responses to drought

From a morphological perspective, plants with smaller stature and reduced leaf area, either by fewer or smaller leaves, tend to have enhanced drought stress tolerance because of their ability to limit water loss and maintain essential physiological functions (Toscano et al., 2019). In our experiment, only minimal differences between the overexpression line and the WT were observed during the stress experiments in terms of height and stem growth.

All plant lines exhibited similar growth rates under control conditions, with no significant differences. Only the stressed HDG11-2 line presented a thicker stem after recovery in the climate chamber experiment. Therefore, the observed drought stress tolerance of the *HDG11*-overexpressing poplars clearly did not arise from reduced growth.

In response to drought, poplars undergo leaf abscission to reduce transpiration and, consequently, water loss when a critical threshold is exceeded. This adaptive response comes at the cost of reduced leaf area, which in turn impairs photosynthetic productivity, highlighting the trade-off between drought tolerance and carbon assimilation in poplars (Larchevêque et al., 2011). Across both experiments, we consistently observed a significantly greater rate of leaf abscission in the WT poplars, which was concomitant with a substantial reduction in biomass production at the end of the climate chamber experiment.

During the 21-day greenhouse experiment, the WT controls produced fewer new leaves than they did at the beginning, whereas the HDG11-2 line maintained a constant leaf-emergence rate under control conditions. The reduced leaf production in the control WT plants over the time of the greenhouse experiment suggests an age-related or nutrient-related decline, further supported by the reduced chlorophyll content after 14 days in the experiment. HD ZIP IV factors are known to regulate epidermal differentiation and meristem activity (Tu et al., 2022). Therefore, *PtaHDG11* overexpression may have contributed to the maintenance of shoot growth capacity in the HDG11-2 line, potentially delaying age- or nutrient-related decline. The lack of a similar effect in the climate chamber experiment may be due to its shorter duration, which may not have been sufficient to induce nutrient deficiency or age-related effects.

### 4.2 Leaf water retention and cell wall-associated regulation

Although all plants were exposed to the same drought treatment, the HDG11-2 line maintained a higher leaf RWC than WT, displaying reduced tissue dehydration which potentially subsequently delayed the onset of stress-induced leaf abscission. A plausible explanation for this phenotype is an altered basal regulation of cell wall remodeling in the transgenic line. XTHs and expansins are key regulators of primary cell wall remodeling, as they modulate wall restructuring, extensibility, and growth-related wall dynamics, and thereby influence turgor-driven cell expansion and tissue mechanics (Tenhaken, 2014; Cosgrove, 2016, 2024). In poplar, *XTH* family genes show pronounced tissue specificity and stress-responsive expression, and their promoters contain multiple stress-related cis-elements, supporting a role in environmentally responsive wall remodeling (Cheng et al., 2021). Moreover, functional evidence from poplar demonstrates that overexpression of the poplar *PagXTH12* in *P*. *alba* × *P*. *glandulosa* enhances drought tolerance, further linking XTH-mediated wall modification to drought adaptation in this genus (Yuan et al., 2024). In parallel, recent work in *A*. *thaliana* showed that expansins, including *AtEXPA15*, are strongly localized in shoot epidermal cell walls and are implicated in cell wall dynamics and drought-related responses, emphasizing that expansin-dependent regulation in leaves is highly tissue- and gene-specific rather than uniformly growth-promoting (Balkova et al., 2025). In our experiment under control conditions, *PtaXTH32*-*like* expression was significantly higher in HDG11-2 than in WT, whereas *PtaEXPA15*-*like* expression was significantly lower. Considering the aforesaid background, the contrasting basal expression of *PtaXTH32*-*like* and *PtaEXPA15*-*like* in HDG11-2 may reflect a modified balance between distinct wall-remodeling processes that affects leaf mechanical properties and thereby the capacity to maintain turgor during water deficit (Cosgrove, 2005; Balkova et al., 2025). Although both genes were strongly downregulated under severe stress, the constitutive differences observed under non-stress conditions suggest that HDG11-2 entered drought stress from a different pre-existing regulatory state than WT, which may have contributed to its improved water retention. This interpretation is particularly intriguing because Xu et al. (2014) showed in *A*. *thaliana* roots that *AtHDG11* directly upregulates several families of cell-wall-loosening genes and binds the promoter of *XTH32* via HD-binding cis-elements. This raises the possibility that *PtaXTH32*-*like*, *PtaEXPA15*-*like*, and potentially other cell wall-remodeling genes, may represent direct or indirect downstream targets of *PtaHDG11*-mediated transcriptional regulation. Taken together, the expression data strongly suggest that altered regulation of cell wall remodeling could have contributed to the water retention that was observed through higher RWC in the HDG11-2 line.

### 4.3 Stomatal regulation and leaf chlorophyll-related responses

Drought performance also depends on how leaf water status is translated into stomatal and chlorophyll-related responses. Decreasing soil moisture generally negatively impairs plant water status, as reflected by reduced RWC, and leads to reduced stomatal conductance as an adaptive mechanism to limit transpirational water loss (Xiong et al., 2022; Pinto et al., 2023). In our experiments, both stressed lines showed reduced stomatal conductance under drought stress. Under severe drought in the climate chamber, HDG11-2 displayed a broad distribution of stomatal conductance values that was positively correlated with leaf RWC. Contrary, the WT maintained uniformly low stomatal conductance regardless of leaf water status, implying a progressive loss of stomatal responsiveness and stress-induced stomatal shutdown in WT. At the transcriptional level, *PtaSDD1* was strongly downregulated under severe drought in both WT and HDG11-2, and notably with an additional significant reduction in the HDG11-2 plants. Overexpression of *SDD1* has been shown to reduce stomatal density in several plant species, including in poplar (*P*. *alba* × *P*. *glandulosa*) (Xia et al., 2024), making *SDD1* a negative regulator of stomatal development. Interestingly, Xia et al. (2024) also show that natural *SDD1* expression in *P*. *alba* × *P*. *glandulosa* increased under osmotic stress, highlighting its role in stress-responsive stomatal patterning but also illustrate genotype-specific differences in *SDD1* expression, when compared to our observations. Further, *PtaSDD1* expression did not differ between WT and HDG11-2 under control conditions, and stomatal density measurements in *in vitro*-grown plants revealed no significant differences between the two lines. Thus, the functional significance of the stronger drought-induced *PtaSDD1* repression in our transgenic HDG11-2 line remains unclear. Given that elevated *SDD1* expression typically reduces stomatal density, conversely, reduced *SDD1* levels would be expected to increase it. Studies on *Quercus robur* and *Populus sp.* report increased stomatal densities with smaller stomata under arid environments. This adaptation enhances stomatal movement sensitivity, thereby accelerating recovery and minimizing trade-offs after drought (Dunlap and Stettler, 2001; Niemczyk et al., 2019; Niemczyk et al., 2024). This suggests that stomatal density is likely strongly dependent on genotype and environmental interactions. It will, therefore, be important in future studies to assess stomatal density and stomatal patterning in relation to *SDD1* gene expression directly under drought conditions in our HDG11-2 line, as the observed positive correlation between RWC and stomatal conductance under drought strongly supports an increase in stomatal movement sensitivity. Overall, since stomatal closure, reflected by stomatal conductance, did not differ consistently between WT and HDG11-2 under stress, altered stomatal closure per se is not the primary factor underlying the enhanced drought tolerance of HDG11-2 but rather stomatal regulation. Surprisingly, stomatal conductance gradually declined in well-watered control plants over the course of the experiments. As these plants were not exposed to soil water deficit, this decrease was not to be caused by drought itself. One possible, though untested, explanation is indirect stress signaling between neighboring plants. Jin et al. (2021) showed that drought-stressed tea plants emit volatile cues that alter stomatal aperture in neighboring plants. Although, to our knowledge, such signaling has not been demonstrated across comparable long-term time scales as our drought stress experiment, these findings raise the possibility that airborne stress cues contributed to the changes in stomatal conductance observed in the control group.

Further, under severe drought, all the plants presented increased chlorophyll content in the upper leaves. A similar response has been reported for poplars under water deficit (Tian et al., 2024). The increased accumulation of chlorophyll in the upper leaf layer may serve to improve light energy absorption or may be a result of carbon redistribution from older to younger leaves to promote photosynthesis (Yao et al., 2024). The response is therefore a general drought acclimation trait rather than a line-specific effect.

### 4.4 Molecular stress markers—proline, MDA, and antioxidant gene expression

Because drought strongly influence cellular stress load, we further assessed molecular physiological and transcriptional markers associated with drought and oxidative stress. To gain insight into the molecular stress response, proline levels, which are known to increase under water stress, were measured. The proline levels did not increase significantly under stress conditions in the *PtaHDG11* overexpression line, whereas they did in the WT plants. The overexpression line did not significantly differ from the basal proline concentrations of the control lines at any stress time point. Interestingly, this observation differs from the results in A*. thaliana*, *F*. *arundinacea*, *O*. *sativa*, *B. oleracea*, and *G*. *hirsutum*, where the overexpression of *AtHDG11* at all times leads to increased proline accumulation under stress and in some cases even under control conditions (Yu et al., 2008; Cao et al., 2009; Yu et al., 2013; Yu et al., 2016; Zhu et al., 2016). Further, the measurement of MDA, a commonly used indirect marker for oxidative stress, showed relatively low levels of MDA in HDG11-2 plants, indicating a reduced level of oxidative stress intensity. Notably, although the MDA levels increased under severe stress conditions in the greenhouse experiment, they rapidly returned to baseline levels after the recovery period. In contrast, the WT plants continued to accumulate MDA during recovery in both experiments, suggesting lingering lipid peroxidation due to inefficient ROS detoxification. This observation implies that HDG11-2 plants may not require proline accumulation, as the cellular stress response appears to be tightly regulated. The transient increase in MDA levels under severe stress conditions may reflect a concomitant rise in ROS, and emerging evidence suggests that MDA itself can act as a signaling molecule to regulate stress responses (Morales and Munné-Bosch, 2019). The higher observed RWC and increased leaf numbers in HDG11-2 suggest greater energy availability for ROS detoxification, enabling better management of oxidative stress and maintaining cellular homeostasis under severe stress conditions (Choudhury et al., 2017). In line with these findings, our gene expression analyses of the antioxidant-related genes *SOD2*-*like* and *CAT2*-*like* revealed significantly higher transcript levels in HDG11-2 under stress conditions in the greenhouse, and also a trend of higher transcript levels under control condition for *PtaSOD2*. Although we cannot conclude that *PtaHDG11* is directly linked to antioxidant gene regulation, the elevated expression of these antioxidant genes may indicate a greater transcriptional readiness of ROS-scavenging pathways in HDG11-2.

### 4.5 Conserved and divergent HDG11-mediated drought responses

Together, these responses indicate that *PtaHDG11* overexpression triggers a broad drought-response in poplar, prompting comparison with previously reported HDG11-mediated effects in herbaceous species. In general, the suite of stress-related responses observed in the *PtaHDG11* transgenic poplars mirrored the patterns reported for *AtHDG11* overexpression in herbaceous species, underscoring the conserved role of this HD ZIP IV TF in conferring drought resilience. Sequence alignment confirmed that the two proteins share approx. 68% overall identity, with > 5% and > 8 % identity of the homeodomain and START domain, respectively, that mediate DNA binding and co factor interaction (Wang et al., 2013). Despite this high degree of similarity, important differences in downstream responses became apparent in the poplar background. Most notably, proline accumulated only in the stressed WT, whereas *AtHDG11* overexpression in herbaceous plants can increase proline accumulation even under control conditions, as mentioned in section 4.4. Proline accumulation is regulated by multiple pathways, including both ABA-dependent and ABA-independent manner (Savouré et al., 1997). To our knowledge, a direct mechanistic link between HDG11 and proline biosynthesis has not been reported. Rather, the increased proline levels reported in herbaceous *AtHDG11* overexpression systems may be associated with enhanced ABA biosynthesis, for example through upregulation of ABA-related genes such as *NCED3*, as reported in Chinese kale (Zhu et al., 2016). Although, neither ABA content nor the expression of ABA biosynthesis genes was assessed in our study, previous work in poplar suggests that drought tolerance can be enhanced without concomitant proline accumulation (Barchet et al., 2014). This pattern is consistent with the response of our HDG11-2 line and suggests that drought adaptation in trees may involve genotype-specific strategies. A second and even more striking example of such context dependence was the complete absence of trichomes in line HDG11-2. This phenotype is consistent with the involvement of HD-ZIP IV TFs in epidermal cell differentiation and trichome development. In *A*. *thaliana*, *AtHDG11* has been implicated in trichome differentiation, whereas the closely related HD-ZIP IV TF *AtGL2* is a well-established positive regulator of trichome formation (Rerie et al., 1994; Nakamura et al., 2006; Lin and Aoyama, 2012; Ahmad et al., 2024). In the affected line, *PtaGL2* expression was significantly upregulated under well-watered conditions. Although this appears contradictory to the trichome-free phenotype, it may reflect the complex regulatory relationship between *HDG11* and *GL2*. Khosla et al. (2014) reported partially redundant functions of *HDG11* and *GL2* in trichome development in *A. thaliana*, indicating that perturbation of HDG11 dosage can affect the regulatory balance of this module. In addition, Ohashi et al. (2002) showed that ectopic *GL2* expression driven by the 35S promoter failed to complement the *gl2* mutant phenotype, whereas entopic expression with the *GL2* promotor could, suggesting that non-native expression can interfere with normal *GL2* function. In this context, the *UBQ10* promoter-driven overexpression used in our study may similarly disturb the spatial and/or temporal regulation of the *HDG11*–*GL2* pathway. Thus, the increased *PtaGL2* transcript abundance may represent a compensatory, but ineffective, response to impaired trichome development rather than a functional restoration of trichome initiation. Accordingly, the trichome-free phenotype of HDG11-2 likely reflects misregulation of the epidermal differentiation program. Recent studies have provided more insight into the role and interaction of HD-ZIP IV TFs in trichome development in various plant species (Cui et al., 2025; Li et al., 2025; Zocca et al., 2025). Given the broad involvement of HD-ZIP IV TFs in epidermal differentiation, root development, and meristem activity (Horstman et al., 2015; Qiu et al., 2022), this finding places *PtaHDG11* within a wider developmental framework in poplar. Our findings are consistent with *PtaHDG11* acting within this broader developmental framework and suggest involvement in epidermal cell differentiation(trichome suppression), although its detailed regulatory network in poplar with respect to trichome development requires further investigation. Notably, altered trichome development has been described for *AtHDG11* in *A*. *thaliana* (Nakamura et al., 2006), where *hdg11* knockout mutants showed the opposite phenotype to our overexpression line, with excess branching of trichomes. Comparable effects of trichome differentiation have, to our knowledge, not been reported in previous heterologous *AtHDG11* overexpression studies in other species, including *P*. × *tomentosa*. Taken together, these findings indicate that, despite the structural conservation of *HDG11*, its downstream regulatory effects are shaped by the species-specific physiological and genetic context and cannot be directly extrapolated from herbaceous systems to woody perennials.

The trichome-free phenotype also raises ecophysiological questions, as trichomes are epidermal structures on plant leaves and shoots that serve as a barrier against pathogens or UV protection and are involved in foliar water uptake (Fambrini and Pugliesi, 2019). Therefore, overexpression of *PtaHDG11* in *P*. *tremula* × *P*. *alba* for field applications might compromise performance under such biotic and abiotic stresses and thus requires further investigation. Because trichome-less HDG11-2 maintained a greater RWC and fewer drought stress symptoms than did the trichome-bearing WT under identical humidity, our observations do not support a major contribution of trichome-mediated foliar water uptake to the drought tolerance observed under the present experimental conditions (Schreel et al., 2020; Li et al., 2023).

Although the present study identified several transcriptional associations linked to *PtaHDG11* overexpression, interpretation of these regulatory relationships should consider that TF activity is ultimately determined at the protein level and may be influenced by post-transcriptional and post-translational regulation. Therefore, transcript abundance alone may not fully reflect *PtaHDG11* regulatory activity under drought stress. Likewise, while the glabrous phenotype coincided with enhanced drought tolerance in HDG11-2, the present study did not directly test whether altered trichome development contributes causally to drought adaptation. Accordingly, the relationships discussed here should be interpreted primarily as transcriptional and phenotypic associations rather than direct mechanistic pathways.

### 4.6 Implications for breeding and forest ecosystem resilience

The findings of this study contribute to our understanding of native *HDG11* function in a woody perennial and point to its potential relevance for the development of drought-resilient tree genotypes. The pronounced improvements in drought stress tolerance, characterized by increased leaf water retention, and subsequent increased dry shoot biomass, identify *PtaHDG11* as a promising target. The use of its native promoter can be incorporated into marker-assisted selection pipelines to accelerate the development of tolerant cultivars for short-rotation coppice and timber production, where water stress can severely curtail biomass yields. The results of the present study should also be extrapolated to other tree species, allowing for the examination of genetic and genomic variation in *HDG11* and its regulatory elements in different tree species. These findings can facilitate the identification of new breeding partners and the development of effective breeding strategies to increase drought stress tolerance in trees. The insights into *HDG11* can guide the selection of drought-tolerant tree genotypes, thereby supporting the development of more resilient and adaptable forest ecosystems.

## Contribution

AF, MF, and TB designed the research. AF performed the research and analyzed the data. AF and TB interpreted the data. AF and TB wrote the manuscript, and all the authors edited the text. MF and TB received the funding.

## Appendix A. Supplementary material

Supplemental Table S1: Primers used for molecular analyses

Supplemental Fig. S1: Transformation vector

Supplemental Fig. S2: Volumetric water content in the greenhouse experiment

Supplemental Fig. S3: Analysis of stomatal density in WT and HDG11-2 poplars

Supplemental Fig. S4: Volumetric water content in the climate chamber experiment under severe stress conditions

## Data Availability Statement

The data of this study are available at the corresponding author upon reasonable request.

## Acknowledgments

We are grateful to Susanne Jelkmann for excellent technical assistance, as well as Alice-Jeannine Sievers, Katrin Groppe, and Vivian Kuhlenkamp for the support within the greenhouse experiment. We thank Sabrina Maninger, Wolfgang Graf, as well as Sabine Benischek and Monika Spauszus for the installation of the drip irrigation system and plant cultivation, respectively. This work was funded by the German Federal Ministry of Food and Agriculture through the funding agency Fachagentur Nachwachsende Rohstoffe (project “TreeEdit”, grant no. 221 NR 5). Open Access funding enabled and organized by Project DEAL.

