## Supplemental Fig. S1 for "Overexpression of *PtaHDG11* enhances drought tolerance and suppresses trichome formation in *Populus tremula* × *Populus alba*"

Fendel A, Fladung M, Bruegmann T*
*) Thünen Institute of Forest Genetics, Grosshansdorf, Germany;

### Supplemental Fig. S1

#### Transformation vector

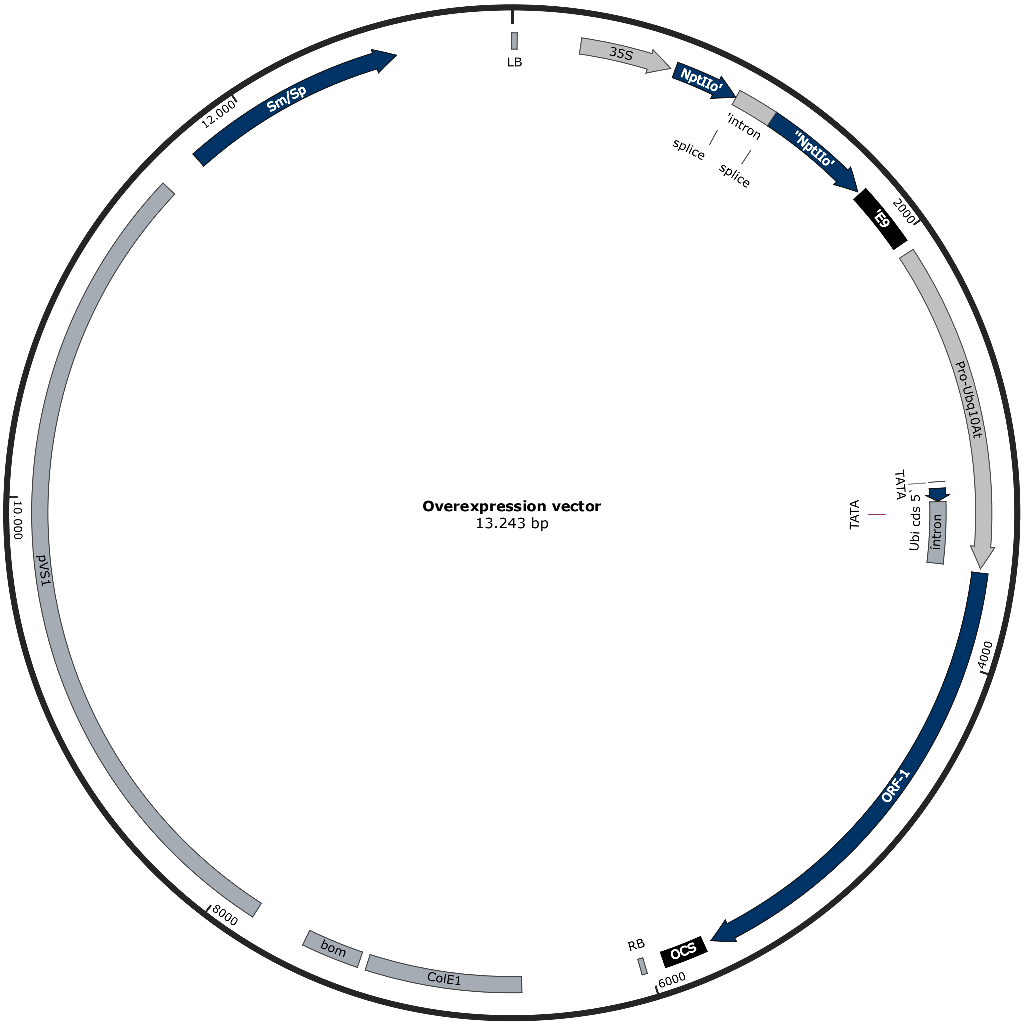


**Fig. S1** Illustration of the overexpression vector used to create the *P. canescens HDG11* overexpression lines. From the left border to the right border of the T-DNA, the vector contains a *Cauliflower Mosaic Virus 35 Promoter* (*CaMV 35S* promoter), a codon optimized *Neomycin Phosphotransferase Two Resistance Gene* (*nptIIo* resistance gene), a *terminator* from *Pisum sativum* (*E9* terminator), an *A.thaliana Ubiquitin promoter* (*AtUbq10* promoter), the open reading frame with the *PtXaAlbH.15G026700* (*PtaHDG11*) CDS, and an *Octopine Synthase terminator* from *Agrobacterium tumefaciens* (OCS terminator). Outside the T-DNA region, the vector contains a *Colicin E1 Origin of Replication* (*ColE1* origin of replication), a *Border of Mobility Region* (*BOM* region), a *Plasmid pVS1 Origin of Replication*, and a *Streptomycin/Spectinomycin resistance gene*
