## Supplemental Fig. S2 for "Overexpression of *PtaHDG11* enhances drought tolerance and suppresses trichome formation in *Populus tremula* × *Populus alba*"

Fendel A, Fladung M, Bruegmann T*
*) Thünen Institute of Forest Genetics, Grosshansdorf, Germany;

### Supplemental Fig. S2

### Volumetric water content in the greenhouse experiment

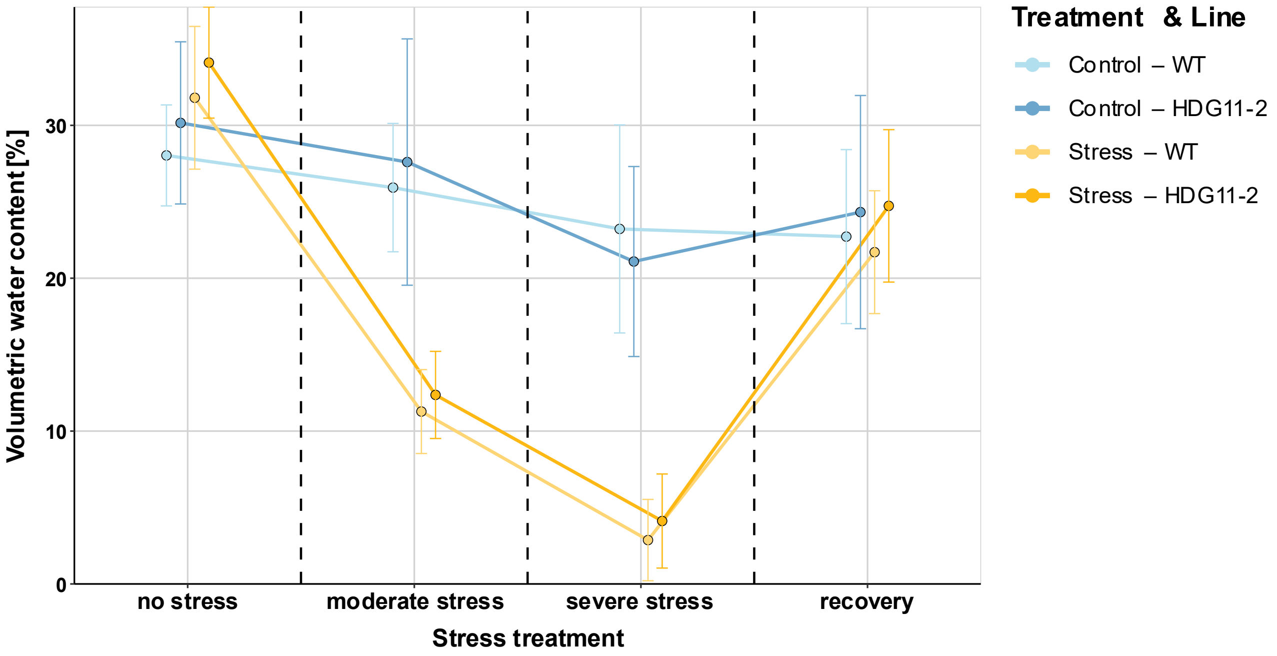


**Fig. S2** VWC of WT and HDG11-2 under drought stress conditions in the greenhouse experiment. The graphs present the means ± SEs ($n_{WT}$ = 7, $n_{HDG11\text{-}2}$ = 4). No significant differences were detected between lines within the same treatment group on a given sampling day
