## Supplemental Fig. S3 for "Overexpression of *PtaHDG11* enhances drought tolerance and suppresses trichome formation in *Populus tremula* × *Populus alba*"

Fendel A, Fladung M, Bruegmann T*
*) Thünen Institute of Forest Genetics, Grosshansdorf, Germany;

### Supplemental Fig. S3

### Analysis of stomatal density in WT and HDG11-2 poplars

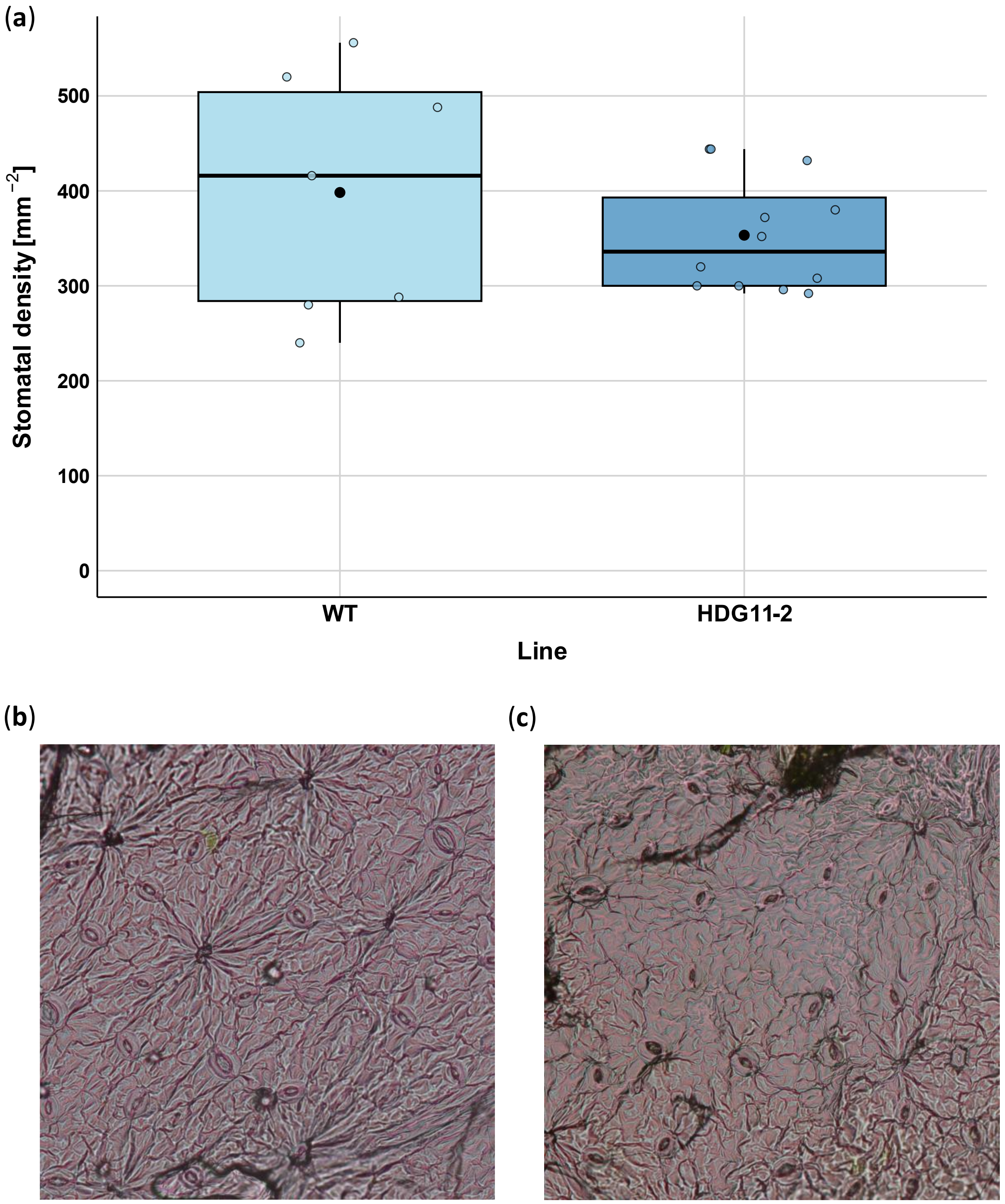


**Fig. S3** Stomatal analysis of *in vitro*-grown plantlets from lines WT and HDG11-2. (a) Stomatal density. The graphs represent the mean ± standard error (SE) of three leaves per line, with three areas analyzed per leaf. No significant differences were detected between the lines, as determined by Student's t-test. Representative sections of stomatal density are shown for (b) WT and (c) HDG11-2
