## Supplemental Fig. S4 for "Overexpression of *PtaHDG11* enhances drought tolerance and suppresses trichome formation in *Populus tremula* × *Populus alba*"

Fendel A, Fladung M, Bruegmann T*
*) Thünen Institute of Forest Genetics, Grosshansdorf, Germany;

### Supplemental Fig. S4

### Volumetric water content in the climate chamber experiment under severe stress conditions

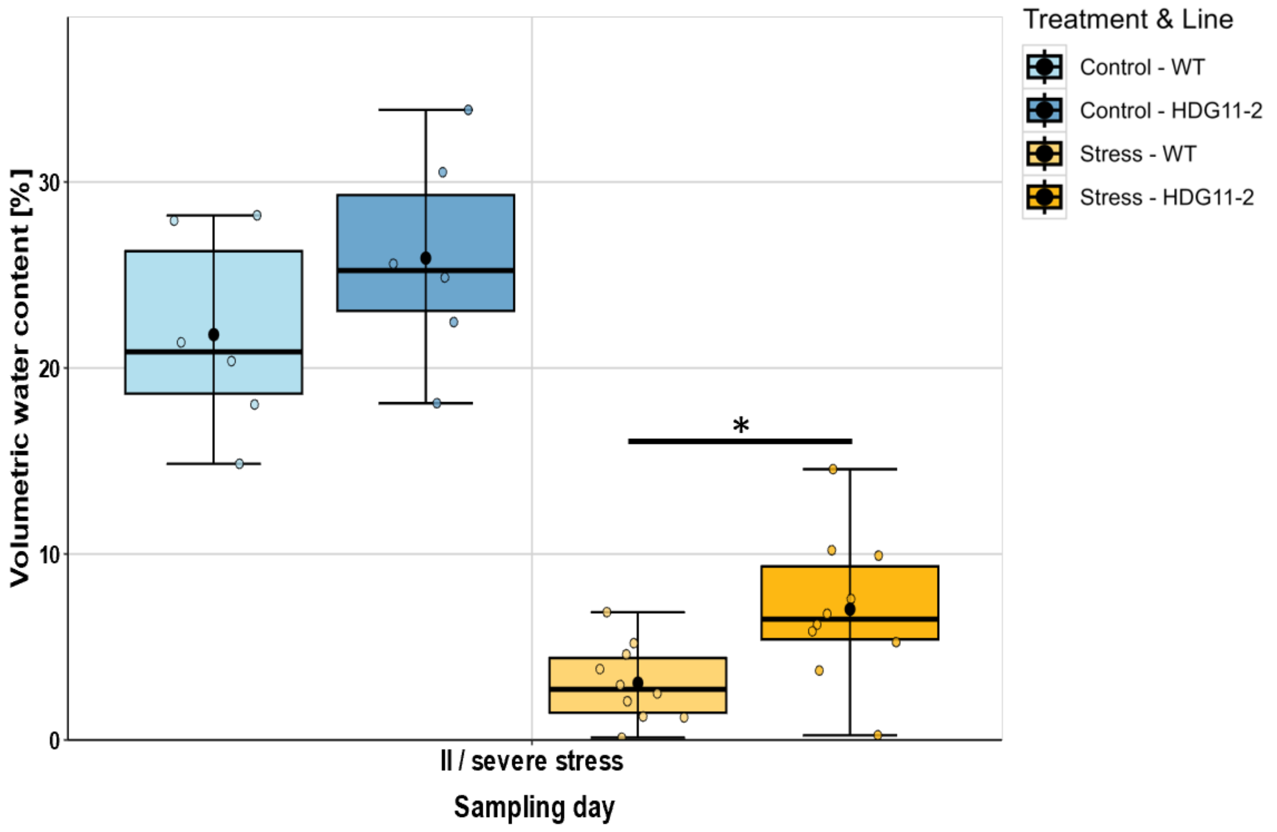


**Fig. S4** VWC of WT and HDG11-2 in response to severe drought stress in the climate chamber experiment. The graphs represent the mean ± SE (control, n = 6; stress, n = 10). Asterisks indicate significant differences between lines within the same treatment, based on Welch-t-test (* $p < 0.05$)
