## Supplemental Table S1 for "Overexpression of *PtaHDG11* enhances drought tolerance and suppresses trichome formation in *Populus tremula* × *Populus alba*"

Overexpression of *PtaHDG11* enhanced drought tolerance and suppressed trichome formation in *Populus tremula* × *Populus alba*
Fendel A, Fladung M, Bruegmann T*
*) Thünen Institute of Forest Genetics, Grosshansdorf, Germany;

### Supplemental Table S1

### Primers used for molecular analyses

| Forward (5'- 3') | Reverse (5'- 3') | Primer binding site(s) | Application |
|---|---|---|---|
| CCAGCCGAGAAGGTCTCCATC | GAACTCGTCAAGCAGGCGGTAG | *nptIIo* | genotyping of transformed antibiotic resistance |
| AGTTTCTAGTTTGTGCGATCG | GGCGTCTCGCATATCTCAT | Pro-*AtUBQ10* (F); Ter-*OCS* (R) | genotyping of transformed target gene *PtaHDG11* |
| ACAGTCTATGCCTCGGGCATC | TCTCGCCTTTCACGTAGTGG | Streptomycin/Spectinomycin resistance gene | test of residual *A. tumefaciens*/plasmids in explants |
| WGTGNAGWANCANAGA | WGTGNAGWANCANAGA | randomly in the genome, 256-fold degenerated | TAIL-PCR |
| / | CGTGAAGTTTCTCATCTAAGCC | T-DNA in the direction of LB | TAIL-PCR |
| / | TCGCCTATAAATACGACGGATCG | T-DNA in the direction of LB | TAIL-PCR |
| / | AGATTTCCCGGACATGAAGCC | T-DNA in the direction of LB | TAIL-PCR |
| CCAGCCGAGAAGGTCTCCATC | GAACTCGTCAAGCAGGCGGTAG | *nptIIo* | Southern Blot probe |
| GTCCACCCTCCATTTGGTCC | GGTCTTTCCTGTGAGCGTCT | *PtaUBQ3/4* | RT-qPCR |
| CCGACCACAGTGGGATGTAC | CCCGGGATGTGAACCATTTGTG | *PtaHDG11* | RT-qPCR |
| GACTACTGCAGATGTAACTGACC | CAATGGCGAAAACTCCTGCC | *PtaSDD1* | RT-qPCR |
| TGCTCCCAAGTGTGCTCATC | TGACCTCCTCATCCCTGTGC | *PtaCAT2-like* | RT-qPCR |
| GAGGATGACCTTGGAAAGGG | ACCAACAACTCCACATGCCAATC | *PtaSOD2-like* | RT-qPCR |
| CCAAGGATTCAACTGGTGTTGC | GCTCCATGTCATTCTTTGTGCC | *PtaGL2* | RT-qPCR |
| GCTAGATAGTACATCAGGAAGTGG | CTCCGGCAGTGTAACCAGG | *XTH32-like* | RT-qPCR |
| CACTTTGATCTCTCTCAGCCTGTC | CCTATAGAAGGGTACCCTGCAGA | *EXPA15-like* | RT-qPCR |
